# Multi-modal profiling of peripheral blood cells across the human lifespan reveals distinct immune cell signatures of aging and longevity

**DOI:** 10.1101/2022.07.06.498968

**Authors:** Tanya T. Karagiannis, Todd W. Dowrey, Carlos Villacorta-Martin, Monty Montano, Eric Reed, Stacy L. Andersen, Thomas T. Perls, Stefano Monti, George J. Murphy, Paola Sebastiani

## Abstract

Age-related changes in immune cell composition and functionality are associated with multimorbidity and mortality. However, many centenarians delay the onset of aging-related disease suggesting the presence of elite immunity that remains highly functional at extreme old age. To identify immune-specific patterns of aging and extreme human longevity, we analyzed novel single cell profiles from the peripheral blood mononuclear cells (PBMCs) of 7 centenarians (mean age 106) and publicly available single cell RNA-sequencing (scRNA-seq) datasets that included an additional 7 centenarians as well as 52 people at younger ages (20-89 years). The analysis confirmed known shifts in the ratio of lymphocytes to myeloid cells, and noncytotoxic to cytotoxic cell distributions with aging, but also identified significant shifts from CD4^+^ T cell to B cell populations in centenarians suggesting a history of exposure to natural and environmental immunogens. Our transcriptional analysis identified cell type signatures *specific* to exceptional longevity that included genes with age-related changes (e.g., increased expression of *STK17A*, a gene known to be involved in DNA damage response) as well as genes expressed uniquely in centenarians’ PBMCs (e.g., *S100A4*, part of the S100 protein family studied in age-related disease and connected to longevity and metabolic regulation). Collectively, these data suggest that centenarians harbor unique, highly functional immune systems that have successfully adapted to a history of insults allowing for the achievement of exceptional longevity.

## Introduction

While a decline in cellular, organismic, and overall functionality is an inexorable outcome of aging, the rate and impact of aging is increasingly recognized to be affected by multiple factors including environment, genetics, and immune history ^3–6^. At a phenotypic level, aging leads to functional abnormalities and alterations in hematopoietic cell populations that prevent a proper immune response and lead to increased susceptibility to infections, cancers, and auto-immune diseases^3–5^. Driven by transcriptional changes and alterations in gene expression, the global immune cell dysfunction generally observed with aging results in distinct shifts in the composition of peripheral immune cell types characterized by a loss of naive B and T cells and an accumulation of memory effector T and B cells ^5,7,8^. In addition, an increase of inflammation-promoting cell populations, such as Natural Killer and myeloid cells (e.g., monocytes) are observed with aging ^5,7,8^, in parallel with gene expression changes in these populations ^8,9^.

At the extreme of the human aging process is extreme longevity (EL), characterized by survival beyond an age reached by less than 1% of a cohort ^10^. EL is often, but not always, associated with a marked delay of disability and in majority (about 60%), common aging-related diseases ^11–13^. Changes in immune cells are considered one of the hallmarks of aging, with growing recognition that the loss of immune competence to control inflammation and rebound from immune stressors is central to the progression of age-limiting morbidities ^14^. Since centenarians–individuals who live to at least 100 years – appear to experience a slower pace of aging ^13,15^, characterizing the repertoire of immune cells of these elite individuals may point to important mechanisms that promote EL.

A recent study using single cell RNA sequencing (scRNA-seq) of peripheral blood mononuclear cells (PBMCs) displayed changes in the distribution of lymphocytes and myeloid cells, and a significant expansion of cytotoxic CD4^+^ T cells in individuals who live to at least 105 years compared to younger individuals ^2^. Hashimoto’s study focused on a cohort of Japanese individuals ^2^, thus it is not clear whether those results generalize to other ethnicities.

Furthermore, the study did not perform an extensive characterization of the transcriptional changes – i.e., changes in expression levels as a function of age and EL – that occur in these cell types.

In this study, we employ a multi-modal approach that combines single-cell transcriptomics with cell-surface protein profiling to characterize both the composition and transcriptional profiles of the peripheral immune system of centenarians. We perform a harmonization of a novel single cell dataset of centenarians with diverse, publicly available peripheral blood scRNA-seq datasets of aging and longevity in an effort to understand the dynamics of circulating immune cell populations throughout the human lifespan, and in particular, in EL.

## Results

### The peripheral immune landscape of centenarians at single cell resolution

Using the 10X Genomics platform for droplet-based Cellular Indexing of Transcriptomes and Epitopes sequencing (CITE-seq) ^16^, we simultaneously profiled the transcriptome-wide expression and the surface-level protein expression of 16,082 PBMCs from seven centenarians (100-119 years of age) and two younger individuals with no known history of familial longevity (20-59 years of age). All nine subjects were of European descent and from the New England Centenarian Study (NECS), with an average sample capture of 1,833 cells **(Supplementary Table S1**). We accounted for technical differences and integrated multiple samples using Harmony ^17^ **(Supplementary Figure 1)**. We then performed Louvain graph-based clustering to group cells into populations of similar expression profiles, and used Uniform Manifold Approximation and Projection (UMAP)^18^ of cell expression profiles to visualize the single cells in a 2-dimensional space.

The identification of major lymphocyte and myeloid cell types was based on 10 immune cell-surface protein expression markers **(Supplementary Figure 2, Supplementary Table S2)**, with subtypes subsequently characterized by transcriptional immune cell signatures previously characterized in human peripheral blood^19^ and fetal liver^20^ **(Supplementary Figure 3-6, Supplementary Table S3)**. This approach identified 11 immune cell types that included major lymphocyte populations: CD4^+^ T cells (CD4TC) with noncytotoxic naive and memory subtypes (nCD4TC, mCD4TC) and cytotoxic subtype (cCD4TC), CD8^+^ T cells (CD8TC) with cytotoxic subtype (cCD8TC), B cells (BC) with naive and memory subtypes (nBC and mBC), and Natural Killer cells (NK) **(Figure 1A, Supplementary Figure 2)**. In addition, we identified major myeloid populations: monocytes with CD14^+^ and CD16^+^ subtypes (M14 and M16) and dendritic cells (DC) with myeloid and plasmacytoid subtypes (mDC and pDC) **(Figure 1A, Supplementary Figure 2)**.

**Figure 1.**
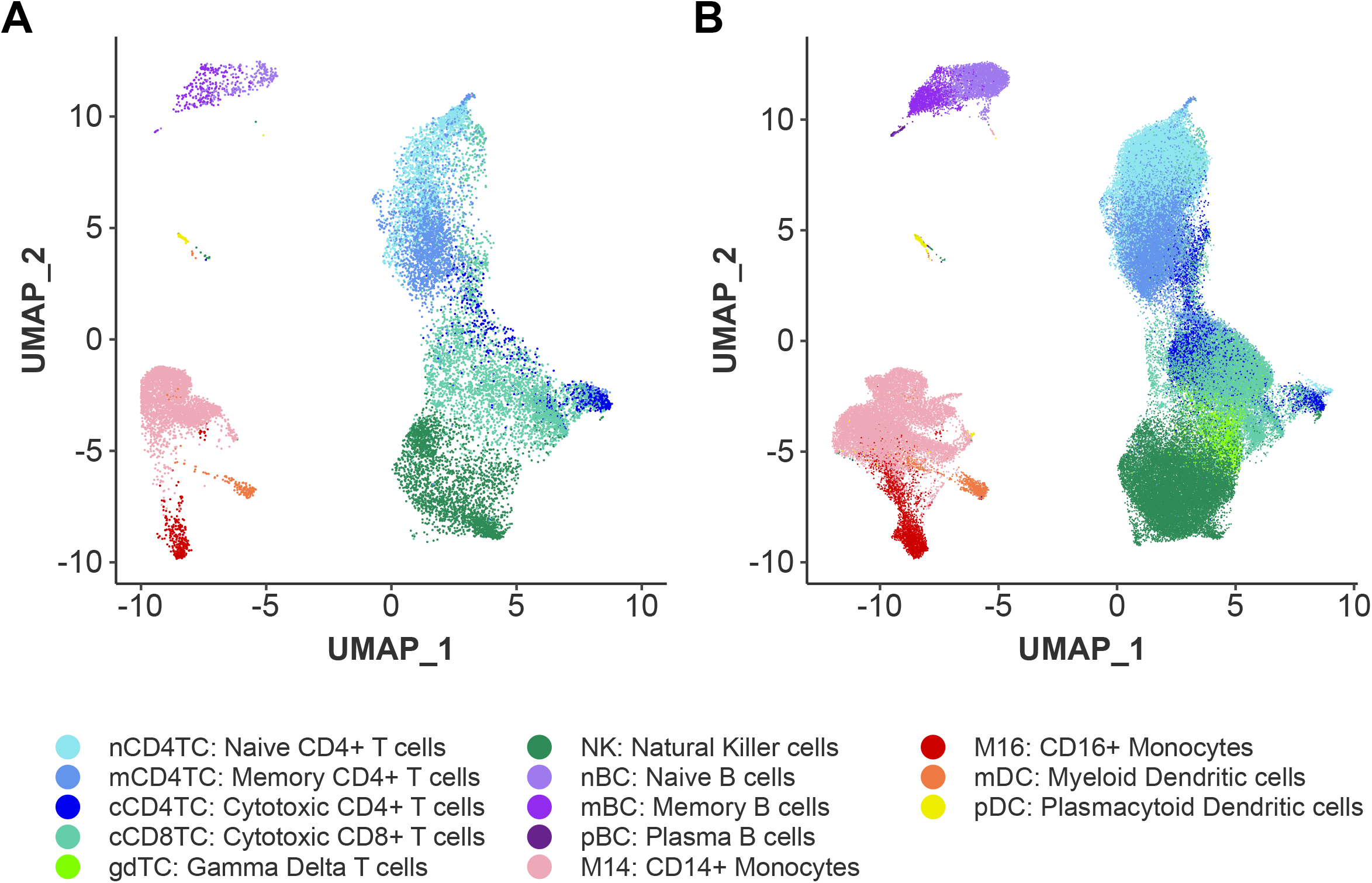
The immune landscape of peripheral blood cells from subjects with extreme longevity at single cell resolution. **A**. UMAP embedding of PBMCs collected from 7 EL individuals from the NECS and 2 younger age controls from the novel CITE-seq dataset, labelled by the identified immune cell subtypes. **B**. UMAP embedding of PBMCs from all 66 subjects representative of the human lifespan from the integrated scRNA-seq datasets (NECS, PNAS, and NATGEN), labelled by immune cell subtypes.

### Centenarians display alterations in immune cell repertoire in comparison to younger age groups

To characterize peripheral immune cell type composition and gene expression profiles across the human lifespan, we integrated our data with two publicly available PBMC datasets of aging and longevity that include subjects of European and Japanese descent ^1,2^ **(Figure 1B, Supplementary Figure 7)**. The integration of these datasets with our novel NECs dataset produced a total of 102,284 cells from 66 individuals across four age groups: 12 subjects of younger age (20-39 years), 26 subjects of middle age (40-59 years), 14 subjects of older age (60-89 years), and 14 EL subjects (100-119 years) **(Supplementary Table S4-S5)**. Technical differences between datasets were accounted for using Harmony ^17^. This analysis identified two additional cell types: plasma B cells (pBC), and gamma delta T cells (gdTC) which were only detected in samples from the Japanese cohort, for a total of 13 cell types across the datasets (**Figure 1B**).

**Figure 2A** displays the observed proportions of the 13 immune cell types in the 66 subjects, stratified by age. Notably, the EL group was characterized by an increase in the proportion of myeloid cells and a reduction in the proportion of lymphocytes: The ratio of myeloid cells to lymphocytes was approximately 13.8/86.2% across the three younger age groups (20-89 years) and shifted to 25.2/74.8% in the EL group **(Figure 2A, Supplementary Table S6)**. This shift in myeloid cells and lymphocytes is an expected trend in aging ^21^. The barplot in **Figure 2A** suggests that the distribution of proportions of the 13 immune cell types becomes more uniform in the EL group. To formalize this observation, we next calculated the cell type diversity statistic of each sample ^22^. This statistic is essentially an entropy-based score that we introduced to summarize the vector of proportions of cell types in a sample. The score is normalized between -1 and 0, with -1 corresponding to the case of a single cell type being present and 0 corresponding to the case when all cell types are present in the exact same proportion and therefore more uniform. The analysis of the 13 immune populations showed a trend towards an increase of the cell type diversity statistic in EL compared to younger age groups, although this difference was not statistically significant (F-test, p-value = 0.7231) **(Supplementary Figure 8, Supplementary Table S7)**. When nCD4TC and mCD4TC were combined as noncytotoxic CD4 T cells, the increase of the cell type diversity statistic in EL compared to younger ages was statistically significant (F-test, p-value = 0.0001875) **(Figure 2B)**. This analysis formalized the observation that the PBMCs of EL subjects comprise more heterogeneity in cell type proportions.

**Figure 2.**
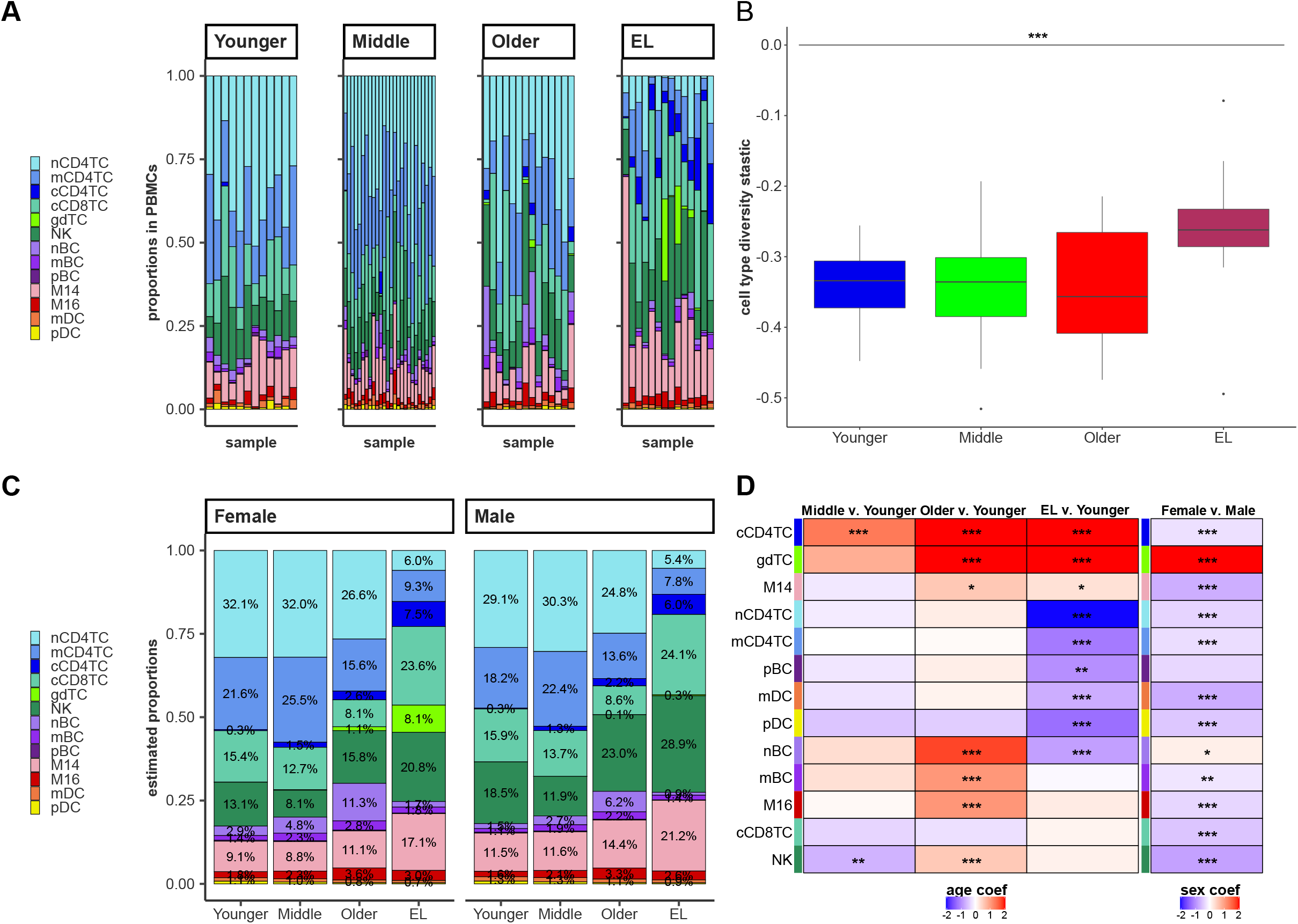
Extreme longevity demonstrates shifts in immune cell repertoire compared to younger control groups. **A**. Bar chart of the relative proportions of the lymphocyte and myeloid immune cell subtypes for each sample across the integrated scRNA-seq datasets : Younger Age, Middle age, Older age, and EL. **B**. Boxplot of cell type diversity statistic calculated on the immune cell subtypes per sample and grouped across age groups, with increased cell type diversity in EL compared to younger age groups although not found to be statistically significant (F-test, p-value =) **C**. Bar chart of the multinomial estimated proportions of the lymphocyte and myeloid immune cell subtypes in each age group, grouped separately for males and females. **D**. Heatmap of the age coefficient comparing Middle, Older, and EL age groups to the Younger age group (right) and heatmap of the sex coefficient for each cell type comparing Females compared to Males. We calculated the Z-statistic and p-value of significance for each coefficient, represented with: *p<0.05, **p<0.01, ***p<0.001.

To estimate the proportions of cell types by age and sex, we next analyzed the observed cell type proportions from all 66 subjects using a Bayesian multinomial logistic regression **(See methods, Supplementary Table S9-S11)**. This analysis produced age and sex specific estimates of each of the 13 cell type proportions across the four age groups (**Figure 2C-D)**. The estimates suggest that there are three main groups of immune cells based on their distributions at different ages and EL: 1) cell types whose proportions increase or decrease monotonically with age and EL (Aging-Related), 2) cell types whose proportions increase or decrease <u>only</u> in the EL group (EL-Specific), 3) cell types whose proportions increase or decrease with age, but these changes do not continue in the EL group (Aging-Specific).

We observed Aging-Related changes (i.e., change in both aging and EL) in cCD4TC, gdTC, and M14 populations. The estimated proportion of cCD4TC increased steadily with increasing age, representing 7.5% of PBMCs in males with EL and 6.0% of PBMCs in females with EL compared to less than 1% of the PBMCs in the younger age groups **(Figure 2C-D)**. This significant change in cCD4TC in centenarians compared to younger controls is consistent with previous findings^2^. We observed a similar aging-related change in the estimated proportions of gdTC and M14 **(Figure 2C-D)**.

Five lymphocyte and myeloid populations were observed to have EL-Specific changes (i.e., change only in EL) and include nCD4TC, mCD4TC, pBC, mDC, and pDC. The lower frequency of nCD4TC and mCD4TC in EL is known ^2^. However, the estimated lower proportion of mDC that decreased to 0.9% of PBMCs in males with EL and 0.7% of PBMCs in females with EL compared to 1.3% in males and 1.1% in females in the younger age group has not been reported. Furthermore, we observed Aging-Specific changes (i.e., change in aging but not EL) in nBC, mBC, and M16 populations. The estimated proportion of nBC significantly increased to 6.2% of PBMCs in males and 11.3% of PBMCs in females in the older age group compared to 1.5% of PBMCs in males and 2.9% of PBMCs in females in the younger age group. However, the estimated proportions of PBMCs that were nBC in EL were 0.9% in males and 1.7% in females.

By examining the proportion of all 13 cell types together, we highlighted major differences in the overall makeup of PBMCs of EL subjects and showed a major shift from innate to adaptive cell types with older age.

### Extreme longevity displays a shift in immune resilience strategy within lymphocyte and myeloid populations compared to younger age groups

To investigate whether the makeup within immune compartments also changed with age and EL and with possible effects on their biological functions and immune resiliency strategies, we next examined the make-up of various immune cell types *within* their myeloid and lymphocyte lineages. To this end, we generated a hierarchy of peripheral immune compartments based on the gene expression profiles of the immune cell types using K2Taxonomer ^23^, and then calculated the average proportions of cell types within each level of the hierarchy (see Methods) **(Supplementary Table S13). Figure 3** displays the results of this analysis and reveals specific changes of the composition of myeloid and lymphocyte compartments in the EL group as compared to younger age groups. We discuss selected results next, with the complete analysis available in **Supplementary Tables S12-25**.

**Figure 3.**
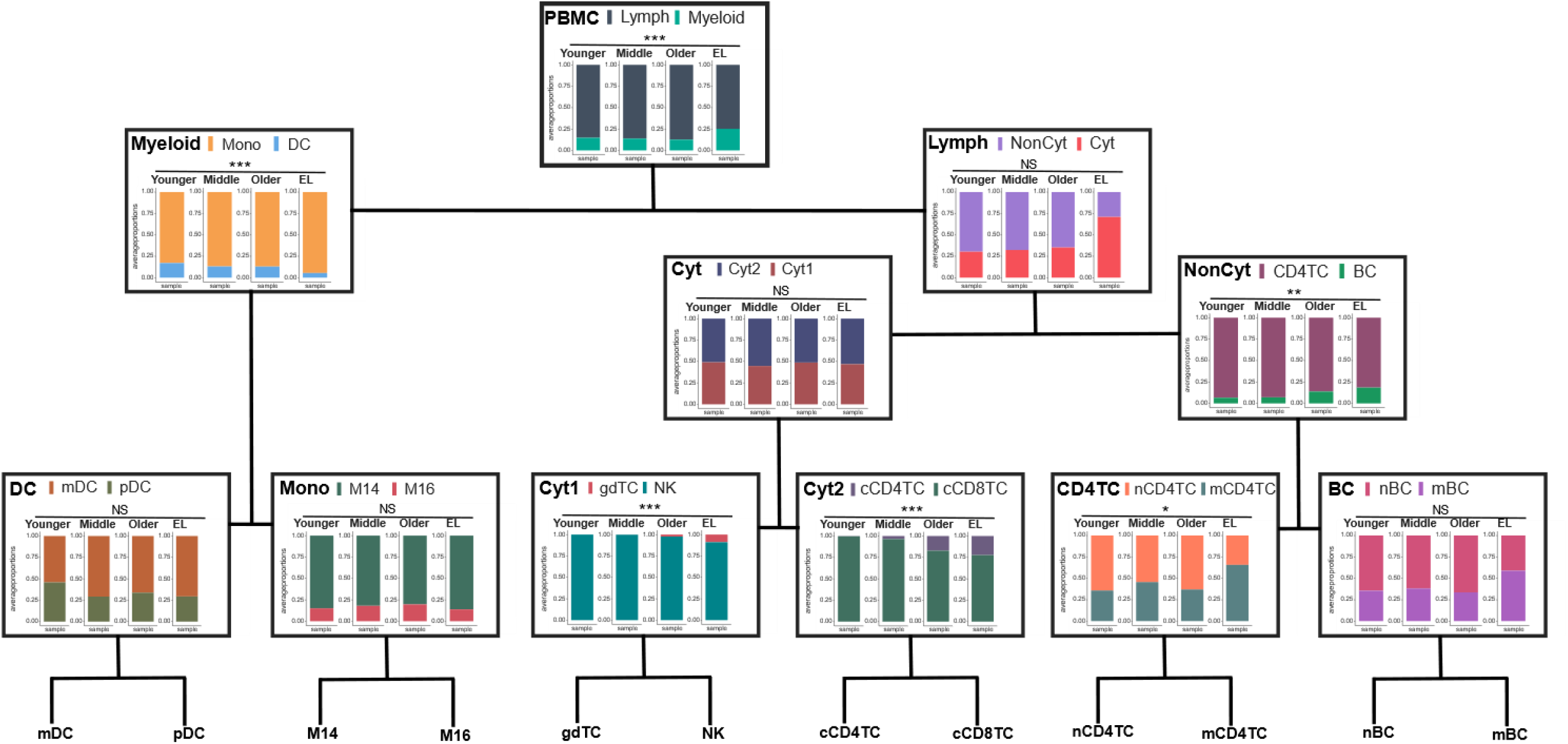
Shift in the immune resilience strategy within lymphocyte and myeloid populations in centenarians. The average immune cell type proportions across the four age groups of the human lifespan along the hierarchy of peripheral immune compartments: PBMC (Myeloid v. Lymph), Myeloid (Mono v. DC), Lymph (NonCyt v. Cyt), DC (mDC v. pDC), Mono (M14 v. M16), NonCyt (CD4TC v. BC), Cyt (Cyt1 v. Cyt2), Cyt1 (gdTC v. NK), Cyt2 (cCD4TC v. cCD8TC), and CD4TC (nCD4TC v. mCD4TC) and BC (nBC v. mBC).

The top of the hierarchy recapitulates the significantly larger proportion of myeloid cells (Myeloid) and smaller proportion of lymphocytes (Lymph) observed in centenarians’ PBMCs discussed earlier. The analysis of the Myeloid compartment (left branch of the tree in **Figure 3**) showed that centenarians’ Myeloid cells were mainly monocytes (Mono) rather than dendritic cells (DC) (EL Mono to DC ratio 94.58%/5.42%) compared to a lower fraction of Mono and higher fraction of DC observed in all other age groups (Mono to DC ratio 85.59%/14.41%). The difference in the two distributions was statistically significant, (F-test of cell type diversity statistic, p-value = 0.0001315) **(Figure 3, Supplementary Figure 9)**. This change in composition of DC has not been reported before ^8,24^. No additional changes were detected within Mono and DC.

The analysis of the lymphocyte compartments (right branch of the tree in **Figure 3**) showed that more than 70% of centenarians’ lymphocytes were Cytotoxic (Cyt: 70.94% and NonCyt: 29.06%) compared to the younger age group (Cyt: 30.30% and NonCyt: 69.70%). This difference in distributions was only borderline statistically significant in our analysis (F-test of cell type diversity statistic, p-value = 0.05032) but consistent with results reported by other investigators^2^. In addition, the hierarchy showed further sub-types of Cyt and NonCyt that had unique composition in centenarians. For example, centenarians’ NonCyt had a significant lower proportion of CD4TC (81.55%) and larger proportion of BC (8.45%) compared to all younger age groups (CD4TC: 90.86% and BC: 9.41%, F-test, p-value = 0.003313). In previous studies, both noncytotoxic CD4TC and BC have been reported to decrease in PBMCs of long-lived individuals ^2^, but this shift between CD4TC to BC has not been previously observed.

The composition of CD4TC in centenarians was characterized by an expansion of mCD4T (65.72%) and a reduction of nCD4TC (34.28%) compared to all younger age groups (mCD4TC: 39.46% and nCD4TC: 60.54%, F-test, p-value = 0.02852). The composition of BC in the EL group had a similar shift from naive (nBC: 41.06%) to memory cells (mBC: 58.94%) but the change did not reach statistical significance (F-test, p-value = 0.7168) **(Figure 3, Supplementary Figure 9)**.

In summary, using the hierarchy of peripheral immune compartments, we identified major shifts in the makeup of centenarian PBMCs compared to those of younger ages within the myeloid and lymphocyte lineages that were obscured globally in terms of all 13 cell types together.

### Centenarians display unique transcriptional profiles associated with Extreme Longevity

The previous two analyses characterized the makeup of centenarian PBMCs globally and within their myeloid and lymphocyte lineages in terms of proportions of the various cell types. We next examined their expression profiles relative to younger age groups. For each cell type, we performed an analysis to discover genes with differential expression as a function of age and/or EL, which identified 99 genes with age- or EL-associated differential expression in at least one cell type (**Figure 4, Supplementary Table S26)**. The number of significantly differentially expressed genes varied by cell types and comparison groups **(Figure 4A)**: on average, the comparison of expression profiles of cell types in the middle *vs*. the younger age group produced a smaller number of differentially expressed genes (i.e., fewer differences) than the comparisons of the older *vs*. younger age and the EL *vs*. younger age group. **Figure 4B** shows clear differential gene expression patterns for all cell type-specific signatures across each age group comparison.

**Figure 4.**
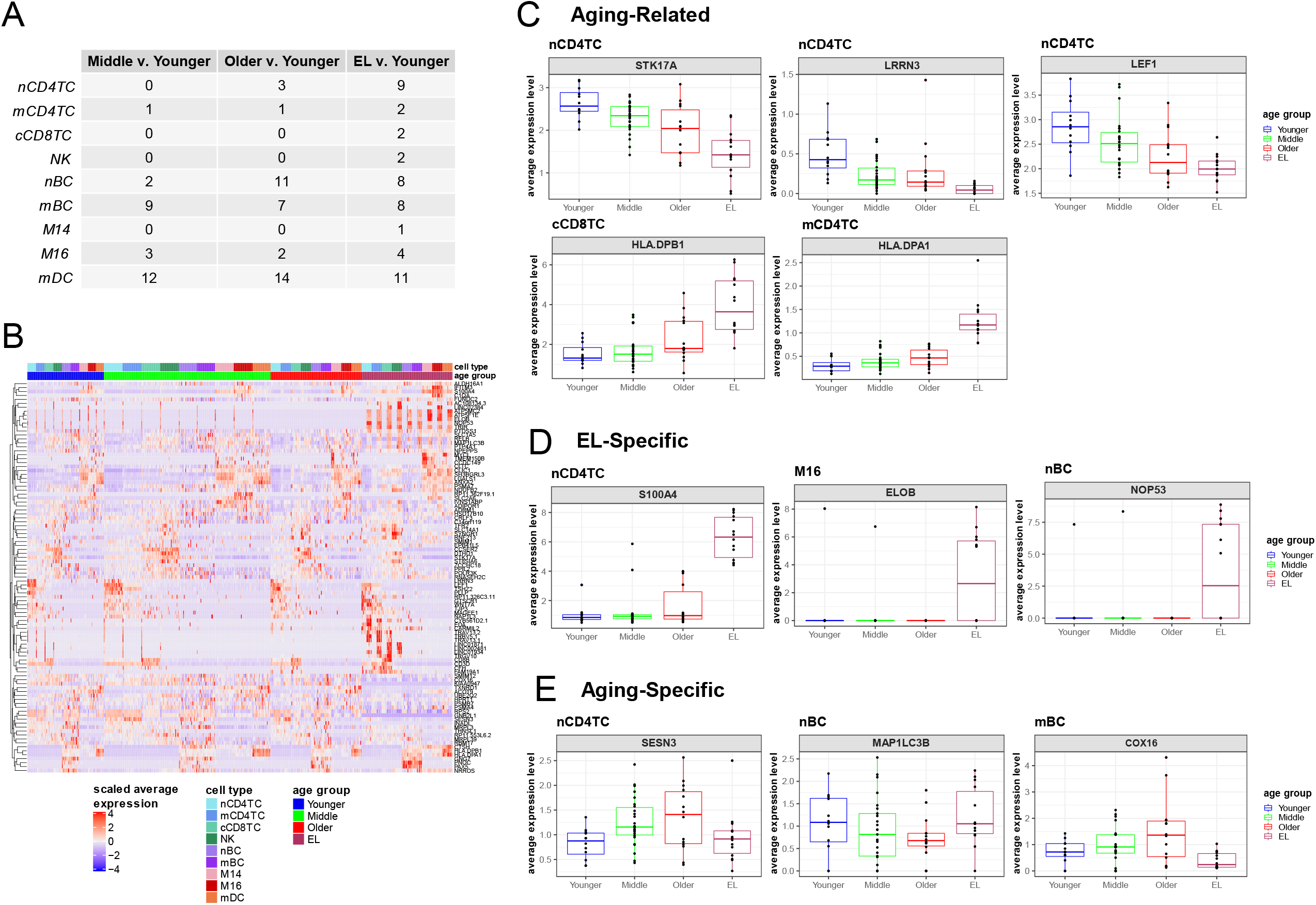
Cell type gene expression changes demonstrate three patterns across the human lifespan. **A**. Table of the number of significant differentially expressed genes across aging comparisons: Middle v. Younger age, Older v. Younger age, EL v. Younger age based on fold change threshold of minimum 10% change and FDR less than 0.05. **B**. Heatmap of scaled average expression per sample of all significant genes across all cell types grouped by age group. **C**. Boxplots of expression levels of specific significant genes in particular cell types demonstrating changes in aging and EL (Aging-Related), **D**. changes only in EL (EL-Specific), and **E**. changes in aging not in EL (Aging-Specific).

Examination of the age-specific expression changes identified three main patterns that are summarized in **Figure 4C-E**, and closely mirror the changes in cell type composition described above. These patterns include 1) genes whose expression increase or decrease monotonically with age and EL (Aging-Related), 2) genes whose expression increase or decrease <u>only</u> in the EL group (EL-Specific), 3) genes whose expression increase or decrease with age, but these changes do not continue in the EL group (Aging-Specific).

We identified 32 genes with the Aging-Related pattern (i.e., change in both aging and EL) of differential expression across the various immune cell types. **Figure 4C** shows selected examples of genes in the first group and **Supplementary Figures 10-18** include more examples. The set includes genes involved in DNA damage response such as serine/threonine-protein kinase 17A (*STK17A)* in nCD4TC **(Figure 4C)**. *STK17A* is a positive regulator of apoptosis and a variant near *STK17A* was previously reported to be associated with longevity in WGS analysis ^25^. Group 1 also included the HLA class II histocompatibility antigen genes *HLA-DPB1* in cCD8TC and the *HLA-DPA1* gene in nCD4TC **(Figure 4C)**. These major histocompatibility genes are involved in antigen presentation and activation of immune response pathways. Furthermore, two genes previously reported to increase in expression with age in peripheral blood were in group 1^25,26^: leucine-rich repeat neuronal protein 3 (*LRRN3*) and lymphoid enhancer-binding factor-1 (*LEF1*) in nCD4TC **(Figure 4C)**.

We identified 8 genes with an EL-Specific pattern (i.e., change only in EL) in several cell types: nCD4TC, cCD8TC, nBC, mBC, and M16 subpopulations **(Figure 4D, Supplementary Figures 10, 14-15, 17). Figure 4D** shows examples of genes that appear to be only expressed in immune cells from the EL group. An example of a gene only expressed in nCD4TC of the EL group is S100 protein-coding gene, *S100A4* **(Figure 4D)**. S100 proteins have been implicated in aging-related diseases such as Alzheimer’s disease as well as longevity ^27,28^. We also identified similar trends in genes involved in transcriptional regulation including Elongin B (*ELOB*, also known as *TCEB2*) in M16, and *NOP53* ribosome biogenesis factor in nBC **(Figure 4D)** and cCD8TC **(Supplementary Figure 12)**.

We identified 11 genes with the Aging-Specific pattern (i.e., change in aging but not in EL), with expression levels that change with age but not in EL across nCD4TC, mCD4TC, nBC, mBC, mDC cell populations. Figure 4E shows examples of 3 genes and additional examples are **in Supplementary Figures 10-11, 14-15, 18**. This set includes genes that respond to oxidative stress including sestrin 3 (*SESN3*) in nCD4TC **(Figure 4E)** and mCD4TC **(Supplementary Figure 11)**, and microtubule-associated proteins 1A/1B chain 3B (*MAP1LC3B*) in nBC, and cytochrome C oxidase assembly factor *COX16* in mBC **(Figure 4E)**. *SESN3* is part of the sestrin family of stress-induced metabolic proteins and is stimulated in response to oxidative stress/damage by *FOXO3*, a transcription factor associated with longevity ^29,30^, while *MAP1LC3B* is involved in autophagy processes in mitochondria of cells and works to reduce oxidative stress ^31^.

In addition to cell type specific expression profiles, we analyzed gene expression aggregated over different cell types (see Methods). This analysis identified a greater number of genes with significantly different expression in the EL v. younger age group comparison (387 genes) compared to middle v. younger age group (0 genes) and older v. younger age group (3 genes) **(Supplementary Figure 19, Supplementary Tables S27)**. Among the 387 significant differential genes identified in EL v. younger age, we found 136 genes over-expressed and 251 under-expressed in the EL group compared to the younger group **(Supplementary Figure 19, Supplementary Tables S27)**. In addition, of the 387 significant genes identified in EL v. younger age, we identified 164 genes previously identified to change in expression with age ^9,32^ including *LEF1* and *LRRN3* ^9,32^ that we also identified at the single cell level as well as *CD28* antigen molecule ^32^.

## Discussion

### Overview of main results

Using a multi-modal, single cell approach, we generated cell composition and transcriptional profiles from the PBMCs of 7 centenarians using CITE-seq. We integrated this novel data set with publicly available scRNA-seq datasets of aging and longevity across the human lifespan to characterize cell type composition and gene expression profiles unique to centenarians. We observed substantial changes in the composition of immune cells with age, including novel changes in myeloid cell types: M14, M16, mDC, and pDC. We also conducted a novel analysis of a data-driven hierarchy of peripheral immune compartments, which revealed previously undetected changes in the composition of T cells and B cells in centenarians. Based on gene expression changes, we identified cell type-specific transcriptional signatures of extreme longevity that include aging-related changes as well as unique gene changes in the immune profiles of centenarians.

### Cell type composition profiles based on the total number of PBMC populations

The peripheral blood immune cell repertoire of individuals is known to change with age ^7,8^. Previous transcriptional studies have shown decreases in lymphocytes and increases in myeloid cells with age ^21^, which we also observed in the peripheral blood of centenarians **(Figure 2)**. However, in addition to these common changes across aging, our analysis identified patterns of immune cell profiles and compositional alterations that are unique to centenarians. We observed expected shifts in the composition of centenarians’ PBMCs from non-cytotoxic (e.g., nCD4TC and mCD4TC) to cytotoxic lymphocytes (e.g., cCD4TC) that have been observed previously in studies of human longevity ^2^. Similarly, the decrease of nBC with aging and longevity has also been reported previously ^7,8^. However, we also discovered novel compositional patterns of extreme old age including aging-related changes (e.g. a significant increase of M14 in older age that continues in the EL group), EL-specific changes (e.g. mDC and pDC display no significant change among the three younger age groups but a unique, significant decrease occurs in EL), and aging-specific changes independent of EL (e.g. a significant increase of M16 in older age that then decreases in the EL age group) **(Figure 2)**. The extent to which these patterns are the drivers or covariates of phenotypic markers of extreme old age remains an open question.

### Cell type composition profiles within peripheral cell compartments

Utilizing the traditional method for characterizing cell type composition profiles based on the total number of PBMC populations, we identified novel compositional changes of extreme old age **(Figure 2)**. However, this analysis provides limited insight into immune cell type change within the lymphocyte and myeloid compartments. We used a novel data-driven approach to create a hierarchy of peripheral immune compartments. The analysis summarized in **Figure 3** shows, for example, that the proportion of lymphocytes in the PBMCs of centenarians decreases compared to younger age groups, but a significant change in composition also occurs. Specifically, centenarians’ lymphocytes are characterized by an almost 50% decrease of NonCyt, which become enriched for BC that are themselves enriched for mBC. Notably, CD4TC and BC have a significant role in the immune system’s response to infection ^33^. BC are associated with the antibody-mediated immune response that triggers a quick response against pathogens, while T lymphocytes such as CD4TC are associated with cell-mediated immunity that develops at a slower rate. Studies have also found crosstalk between BC and CD4TC, showing their co-dependence in protective immune response ^33^. The shift from CD4TC to BC suggests that centenarians develop a more immediate immune response to infections. We also observed a significant shift from naive to memory subtypes within CD4 TC and, to an extent, within BC suggesting that centenarians did not escape infection but experienced a greater exposure to infections and were able to develop robust responses to them. Previous studies have also shown associations between the capacity to control inflammation and preserving immunocompetence with longevity as an immune resilience phenotype ^34–36^. It is possible that the unique make up of immune cells we observed in centenarians may represent an adaptation of their immune system or a compensatory mechanism to the loss of key immune cell types. Interestingly, in almost all compartments displayed in **Figure 3**, the EL group was characterized by a lesser skewed distribution of cell types compared to younger age groups. A more heterogeneous distribution of immune cells may be the driver of their immune resiliency. Compositional heterogeneity may reflect immunocompetence as a dynamic balance or homeostasis ^36^. Reciprocally, immune imbalance is often characteristic of suboptimal responses to infections (e.g., COVID-19 ^37^) or compensation by less effective or exhausted cell mediators (e.g., NK cells ^38^) that compromise health span.

### Gene expression profiles

Our analysis identified three patterns of age-related changes: monotonic changes across the lifespan, age-related changes that are absent in the EL group, and changes that are unique to centenarians. Interestingly, we noticed similar patterns in the serum proteome of centenarians ^39^. By comparing the serum proteome of centenarians to septuagenarians, we discovered aging related protein signatures as well as a protein signature that was unique to centenarians. We also discovered an age-specific protein signature that was not extended as expected in centenarians. Some of the genes with differential expression in the EL group have been linked to aging and longevity studies. For example, the expression of a variant of *STK17A* that we found associated with age in nCD4T (Figure 4C) was higher in centenarians ^25^. *STK17A* is involved in DNA damage response, positive regulation of apoptosis, and mitochondrial and metabolic regulation of reactive oxygen species (ROS). This association is consistent with results from previous studies that correlated DNA damage repair mechanisms to aging and longevity ^25,40,41^. *S100A4*, part of the S100 family of calcium-binding proteins, showed an EL specific change in naive CD4 T cells (Figure 4D). S100 proteins such as *S100A13* have been implicated in their role in longevity, including an association with *APOE* genotypes in centenarians ^28^. In addition, the S100 family of proteins are associated with inflammatory pathways in the brain connected to aging-related diseases such as Alzheimer’s disease ^27^. A recent study discovered an S100 protein to be a critical regulator of hematopoietic stem cell renewal through mitochondrial metabolic regulation and function ^42^. In rats, recombinant *S100A4* demonstrates an anti-apoptotic function in response to oxidative stress injury ^43^. Other transcriptional signatures that we identified in our analysis, such as *SESN3* and *MAP1LC3B* (Figure 4E), are involved in DNA damage response and mitochondrial and metabolic regulation activated in response to oxidative stress ^29,30^. Sestrins such as *SESN3* are highly conserved stress inducible proteins that protect the immune system in response to DNA damage and oxidative stress ^30^. More specifically, the induction of *SESN3* in response to oxidative damage is activated by *FOXO3* ^30^, a transcription factor associated with longevity ^29^. *MAP1LC3B* is involved in autophagy processes in mitochondria of cells and works to reduce oxidative stress ^31^. In the immune system, mitochondrial regulation plays a role in immune cell transcription and activation ^44,45^ and may promote longevity ^46^. In addition, the decline of mitochondrial quality and activity is associated with aging and senescence ^44,45^. The connection to mitochondrial and metabolic regulation suggests that centenarians may have changes that occur in mitochondrial and metabolic regulation and function and should be further investigated. We note that our signatures of aging derived from “pseudo-bulk” data closely match previously published signatures of aging derived from bulk data ^9,32^. The difference between the bulk and single-cell signatures on the other hand, point to the fact that much of the observed differential expression is driven by differences in cell composition rather than by differences in within-cell type expression.

### Limitations and Future Directions

This study has several limitations, particularly the cross-sectional nature of the data and the small sample size. We integrated our data with multiple single cell datasets to increase sample size, but we intentionally adopted a conservative approach to identify cell type specific signatures across age groups. Larger studies of centenarians will be needed to detect robust transcriptional changes that characterize EL. Although we identified a small set of transcriptional signatures, we were still able to identify patterns of EL that have been discovered in previous studies including EL specific changes. In addition, the compositional and gene expression changes that we observed in centenarians displayed not only EL specific changes but also age-related changes. How EL differs from regular aging remains unclear, and more investigation and future studies will be required to elucidate this difference and investigate the mechanisms behind the patterns observed in extreme old age. Access to the peripheral blood of centenarian offspring and studying longitudinal changes in PBMC populations may help to better define immunocompetence causal drivers of the beneficial health outcome observed in EL.

### Conclusion

Overall, these findings display age-related changes in composition and transcription in both lymphocyte and myeloid cell types that collectively reflect immunocompetent profiles that may in part account for centenarians’ ability to reach extreme ages. The extent to which some of the unique compositional and transcriptional patterns we identified in centenarians’ PBMCs are the drivers or markers of extreme old age remains an open question. To our knowledge, this is the first study to define cell compositional and transcriptional signatures of EL across immune cell types in peripheral blood. This study provides a foundation and resource to explore immune resilience mechanisms engaged in exceptional longevity.

## Methods

### Experimental Procedure

#### Human blood samples

Centenarians were enrolled in North America in 2019. The study was approved by the Boston Medical Center and Boston University Medical Campus IRB and all participants provided written informed consent.

#### Processing of blood samples

For each centenarian and control individual involved in this study, 8 mLs of peripheral blood was drawn into each of two BD Vacutainer Cell Preparation Tubes with sodium citrate (BD Biosciences catalog #362760). The tubes were centrifuged at 1,800 x g for 30 minutes at room temperature (RT). The buffy coat containing peripheral blood mononuclear cells (PBMCs) was isolated and transferred into a sterile 15 mL conical centrifuge tube. The PBMC sample was brought to 10 mLs with sterile Dulbecco’s phosphate buffered saline (DPBS, Invitrogen catalog #14190-144) and centrifuged at 300 x g for 15 min at RT. The supernatant was aspirated and the pellet resuspended in 10 mL sterile DPBS and a cell count performed via hemocytometer. The sample was centrifuged at 300 x g for 10 min at RT and the supernatant aspirated. The pellet was resuspended in chilled (4°C) resuspension medium (40% fetal bovine serum (FBS) hyclone defined, Cytiva catalog #SH30070.03 in Iscove’s Modified Dulbecco’s Medium (IMDM), catalog #12440053) to achieve a cell concentration of 4 × 10^6 cells/ mL. An equal amount of chilled 2X freezing medium (30% Dimethyl Sulfoxide (DMSO), Sigma catalog #D2650 in IMDM/ 40% FBS) was added to achieve a cell concentration of 2 × 10^6 cells / mL. This mixture was then aliquoted into 1.2 mL cryovials (Corning catalog #430487) at 1 mL / vial. These vials were then brought to -80°C before being transferred to a -150°C deep freezer.

#### scRNA-seq of PBMCs of centenarians

PBMC samples (2 × 10^6 cells/sample) from the centenarian cohort were thawed rapidly and mixed with 15 mL StemSpan medium (CAT#) with L-glutamine (1:1000 conc…). These samples were then brought to 50 mL with sort buffer (2% Bovine Serum Albumin (BSA), Millipore Sigma catalog #EM-2930 in DPBS) and centrifuged for 5 min at 400 x g at RT. The supernatant was aspirated, and the pellet of PBMCs resuspended in 30 mL sort buffer and centrifuged for 5 min at 400 x g at RT. The pellet was resuspended in 2 mL StemSpan medium (+L-glutamine) and filtered through a 40 uM filter. The samples were then incubated in this medium for 1 hour at 37°C/5% CO2. Following this incubation, the samples were centrifuged for 5 min at 400 rcf at RT and the supernatant aspirated. Each pellet was then resuspended in 50 uL labeling buffer (1% BSA in PBS) and 5 uL Human TruStain FcX() was added. The samples were incubated at 4°C for 10 minutes. During this incubation, the TotalSeq-C antibody pool containing 1 ug of each antibody was prepared and centrifuged for 10 min at 14,000 x g at 4°C. The supernatant was then transferred and used as the antibody mix for each sample. 20 uL of the antibody mix was added to each sample and the samples were brought to 100 uL with labeling buffer. The samples were incubated for 30 minutes at 4°C. Following this incubation, the samples were washed with 1.3 mL labeling buffer and centrifuged at 400 x g for 5 minutes at RT. This washing step was repeated for a total of 3 washes. The pellets were then resuspended in 500 uL labeling buffer with calcein blue AM(1:1000) Fisher Scientific, catalog #C1429). 1 × 10^5 calcein blue positive cells (live cells) were then sorted per sample. The sorted samples were centrifuged for 5 min at 400 rcf at room temperature and resuspended in 120 uL resuspension buffer (0.04% BSA in DPBS). The samples were counted via hemocytometer and diluted to 600 cells/uL with resuspension buffer.

### Single cell analysis

#### New England Centenarian dataset

##### CITE-seq and CellRanger Preprocessing

Cellular Indexing and epitopes sequencing (CITE-seq) was performed on the 7 centenarians and 2 younger age controls using a commercial droplet-based platform (10x Chromium). We constructed 5’ gene expression libraries (GEX), as well as surface protein libraries (antibody derived tags, ADT) following the manufacturer’s user guide. These libraries were sequenced on two runs of an Illumina NextSeq 2000 instrument generating 438 and 535 million reads respectively.

Raw sequencing files were converted to fastq and demultiplexed using bcl2fastq v.2.20 and Cellranger v.3.0.2. Counts for the expression and antibody capture libraries were derived by simultaneously mapping the respective fastq files to the human genome (GRCh38) and to the feature reference of the TotalSeq-C antibodies used using the corresponding parameters in cellranger count (v.3.0.2). This pipeline includes the alignment, barcode and UMI counting.

##### Filtering, PCA Analysis, Batch Correction, and Clustering

After processing the samples through CellRanger, we performed filtering, normalization, and principal component analysis using Seurat v.3 ^47^. First, we performed quality control steps based on the number of genes and UMIs detected per cell, and percent of mitochondrial genes expressed per cell. To remove poor quality cells with low RNA content, we removed cells with less than 200 genes detected. To filter out outlier cells and doublets, we filtered out cells with greater than 3,000 detected genes, as well as cells with greater than 15,000 UMIs. To account for cells that are damaged or dying, we removed cells with greater than 15 percent mitochondrial counts expressed. After filtering, we normalized the RNA-level expression data for each cell to compare gene expression between sample cells; Gene counts for each cell were normalized by total expression, multiplied by a scale factor of 10,000 and transformed to a log2 scale. We normalized the protein-level expression data by applying a centered log ratio (CLR) transformation for each cell to account for differences in total protein ADT counts that make up each cell.

Further downstream analyses were performed on the RNA-level expression data. PCA based on the top 2000 highly variable genes was performed for dimensionality reduction, and the top 20 significant PCs were selected that explain the most variability in the data. The top significant PCs were used as an input for clustering the cells and for nonlinear dimension methods mentioned below to identify populations of cells with similar expression profiles.

To account for technical variations between samples from different experimental batches, we corrected the PCA embeddings using the Harmony algorithm ^17^, a method that iteratively clusters and corrects the PC coordinates to adjust for batch specific effects. We assessed the integration of these datasets by employing PCA visualizations of batches of cells and calculating the average silhouette width (ASW) score ^48,49^ for each cell type population based on the top 20 principal components before correction and the top 20 harmony components after batch correction reported with T-statistic and p-value significance threshold of 0.05.

We clustered cells based on graph-based methods (SNN and Louvain community detection method) using the top 20 Harmony-adjusted components and used the Unifold Manifold Approximation and Projection (UMAP) algorithm^18^ to visualize clusters of cells and other known annotations.

##### Identification and classification of cell types

We used a multi-modal approach to identify immune subpopulations in the NECS dataset. First, we used the 10 cell-surface protein immune cell marker panel of expression to identify main immune cell types. We then further partitioned the main immune cell types into immune subtypes using graph-based clustering and based on the expression of immune cell type signatures from literature ^19,20^. The average expression score of each signature was calculated for a single cell by calculating the average scaled expression of all genes within a signature, with the scaling based on the expression of a control set of genes (AddModuleScore function in Seurat ^47^), and by taking the absolute value of the average scaled expression to compare scores across signatures within a cell population.

#### Publicly available datasets

##### Data collection, filtering, PCA analysis, and clustering

We downloaded the raw UMI matrix for the scRNA-seq dataset of PBMCs from 45 younger age controls of European descent ^1^, which we will refer to as NATGEN. We also downloaded the raw UMI matrix for the scRNA-seq dataset of PBMCS from 7 supercentenarians and 5 younger age controls of Japanese descent ^2^, which we will refer to as PNAS. For both PBMC datasets, we performed all downstream processing including filtering, normalization, and scaling of data using the Seurat v.3 ^47^. For both datasets, we performed quality control steps based on the number of genes and UMIs detected per cell, and percent of mitochondrial genes expressed per cell. For the NATGEN dataset, we filtered cells based on similar thresholds from the original manuscript ^1^ with the exception of filtering the number of UMIs per cell greater than 3,500 to remove outlier and doublet cells and filtering the percent of mitochondrial genes expressed greater than 5 percent to remove damaged or dying cells. After filtering, the dataset included UMI counts for 25,000 cells. For the PNAS dataset, we filtered cells as previously published ^2^. After filtering the datasets, we normalized the expression levels of each cell to compare gene expression between sample cells; gene counts for each cell were normalized by total expression, multiplied by a scale factor of 10,000 and transformed to a log2 scale. We then performed PCA analysis based on the top 2,000 highly variable genes detected and clustered cells based on graph-based methods (SNN and Louvain community detection method) ^47^ based on the top significant PCs for each data set implemented in Seurat. We used the UMAP algorithm ^18^ to visualize the clusters of single cells and other known annotations.

##### Subpopulation Identification and Harmonization

To define the set of consensus immune cell types across regular aging and longevity, we first identified subpopulations of each cell type using immune cell type signatures from literature ^19,20^. The average expression score of each signature was calculated for each single cell across datasets as described for the NECS dataset using Seurat. In addition, we compared the expression of canonical gene markers of the immune populations identified for comparison ^50^. After identifying subpopulations in each dataset, we then integrated these scRNA-seq datasets of PBMCs by correcting the PCA embeddings using the Harmony algorthim^17^ and assessed batch correction using average silhouette width (ASW) score ^48,49^ for each cell type population based on the top 20 principal components before correction and the top 20 harmony components after batch correction reported with T-statistic and p-value significance threshold of 0.05.

#### Overall cell type composition analysis

We compared the overall cell type composition differences across samples and age groups by calculating the cell type diversity statistic for each sample, a normalized entropy-based method borrowed from the analyses of microbiota data ^22^. The cell type diversity statistic Es is represented as the measure of entropy based on the cell type proportions pi for the sample s normalized based on the total number of cell types k:

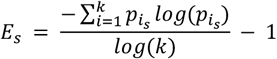

A sample with more uniformity in cell type abundances/proportions will result in having a higher cell type diversity statistic compared to a sample with cell type abundances/proportions that are skewed towards specific cell types will have a lower diversity statistic. We performed ANOVA with p-value of 0.05 to assess differences between age groups.

#### Cell type specific composition analysis

To investigate cell type specific differences across age groups, we applied a Bayesian multinomial logistic regression model to the cell type abundances:

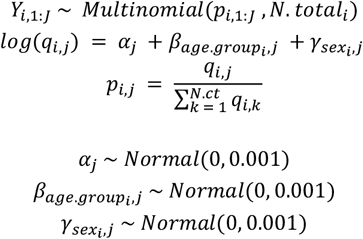

where *Y*_*i*,1*:J*_ represents the abundances of cell type 1: *J* for sample *i* that are modeled using a Multinomial distribution with probabilities 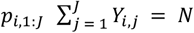 for all sample *i* and 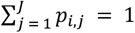. The probabilities *p*_*i*,1*:J*_ depend on age and sex through the function *log(q*_*i,j*_). We chose a reference for each age group (younger age) and sex (male).

The model was estimated using Markov Chain Monte Carlo (MCMC) sampling using rjags, the R package for JAGS ^51^. We ran parameter inference for all coefficients for group level probabilities of composition (*P*_*i,j*_) and the age group and sex effect coefficients (*β, γ*) using 1,000 iterations with 500 iterations for burn-in.

We obtained the group level composition probabilities and 95 percent credible interval for males and female subjects for younger, middle, older, and EL age groups across all immune cell types. In addition, to assess the significance of the effect of age and sex in each cell type, we calculated the z-score (Z) for parameters *β* and *γ* based on the mean estimate and standard error of the posterior distribution. we subsequently calculated the two-sided p-value based on the standard normal distribution: 2Φ(−|Z|)) where Φ is the standard normal cumulative distribution function. We calculated the adjusted p-value based on the Benjamin and Hochberg correction for multiple testing across all coefficients tested.

#### Analysis of the hierarchy of peripheral immune compartments

We investigated differences in immune cell composition between age groups across multiple molecularly derived cell type subgroups comprising peripheral immune compartments. To estimate this hierarchy, we utilized K2Taxonomer (v1.0.5) ^23^, which performs top-down partitioning of cell types based on the relative similarity of their transcriptomic profiles. Prior to running K2Taxonomer, we performed several further data processing steps. First, plasma cells were removed from this analysis because they were present in fewer than 10 subjects. Next, for each subject we aggregated single-cell profiles of each cell type into a “pseudo-bulk” profile by summing the counts of each gene, followed by normalization to log2(counts-per-million). The resulting data set included 515 total profiles across 66 subjects and 12 cell types. The number of subjects for which each of these 12 cell types were identified in each of the four batches is given in Supplementary Table S12. Next, we removed lowly expressed genes, which failed to reach 2 counts-per-million in at least 2 profiles across all batches, which left 12,354 remaining genes. Finally, we performed batch correction on these data using ComBat (v3.40.0) ^52^.

To compare the cell composition between age groups, we performed cell type diversity statistic-based analyses, independently, for each subset of cell types comprising peripheral immune compartments estimated by K2Taxonomer. Statistically significant differences in mean cell type diversity statistics were assessed using ANOVA with p-value threshold of 0.05.

#### Cell type specific differential gene expression analysis

To investigate the cell type specific differences across age groups, we created and applied a Bayesian normally distributed mixed effects model with the rjags R package ^51^ to compare gene expression changes between middle, older, and EL age groups in reference to the younger age group. For each cell type, we first filtered genes to keep genes with expression in at least fifty percent of the smallest cell type population. We then performed differential gene expression analysis across the four age groups (younger, middle, older age groups) using the Bayesian mixed effects model where for each cell (i):

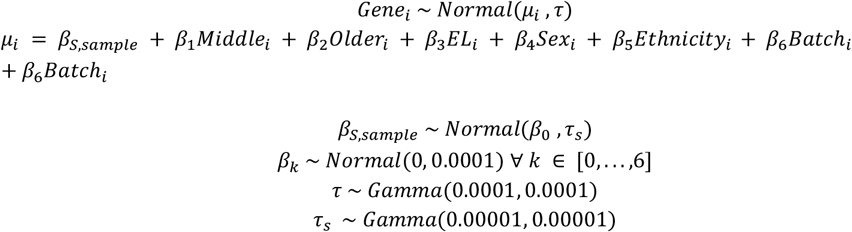

where *Gene*_*i*_ is the log-normalized expression of a gene for each cell *i* in the cell type; *β*_1_, *β*_2_, *β*_3_, *β*_4_, *β*_5_, *β*_6_, *β*_0_ are model parameters. The model is adjusted by fixed covariates sex, batch, and ethnicity, as well as a random effect based on samples to account for differences in cell abundances between samples within groups. Using this model, we monitored the age-dependent coefficients (β_1_, β_2_, β_3_) across 10,000 MCMC iterations with 2,500 burn-in iterations to obtain the log fold change (logFC) based on age group. Then, we calculated the z-score (Z) for the age-dependent parameter based on the mean estimate and standard error of the posterior distribution. We subsequently calculated the two-sided p-value based on the standard normal distribution: 2*Φ(-|Z|)) where Φ is the standard normal cumulative distribution function. We calculated the FDR based on the Benjamin and Hochberg correction for multiple testing across all genes tested. Significant differential genes were selected based on a significance of FDR < 0.05 and fold change cutoff of 10 percent (logFC < |log2(1.1)|).

#### Bulk level differential gene expression analysis

We performed differential gene expression analysis at the bulk level between age groups using DESeq2 ^53^. We filtered genes in the single cell data to keep genes with expression in at least 50% of the smallest cell type population. We then aggregated the raw counts per sample and ran DESeq2 to perform normalization and fit a negative binomial generalized linear model with covariates age group, sex, batch, and ethnicity. A Wald test was subsequently performed on each age group coefficient (i.e., Middle, Older, and EL v. Younger age) with log fold changes, Wald p-values, and adjusted p-values (FDR) reported. Differentially expressed genes at the bulk level were evaluated according to log fold change greater than log2(1.5) and FDR < 0.05.

## Data Availability

The data that support these findings are publicly available and were accessed from several repositories. NATGEN single cell expression data and subject level data were publicly available as referenced in ^1^: https://molgenis58.target.rug.nl/scrna-seq/. PNAS single cell expression data and subject level data was available as referenced in ^2^: http://gerg.gsc.riken.jp/SC2018/. NECS will be available from Synapse (https://adknowledgeportal.synapse.org/Explore/Projects/DetailsPage?Grant%20Number=UH2 <u>AG064704</u>).

## Code Availability

All scripts to reproduce analyses and figures reported in this paper are available on github (https://github.com/Integrative-Longevity-Omics/sc_pbmc_centenarians).

## Acknowledgements

TK, SM, PS, GM, SA, TP are supported by NIH-NIA UH2AG064704. MM and PS are supported by NIH NIA Pepper center: P30 AG031679-10.

