## Supplementary Figure for "Multi-modal profiling of peripheral blood cells across the human lifespan reveals distinct immune cell signatures of aging and longevity"

### Supplementary Materials

### Supplementary Figures

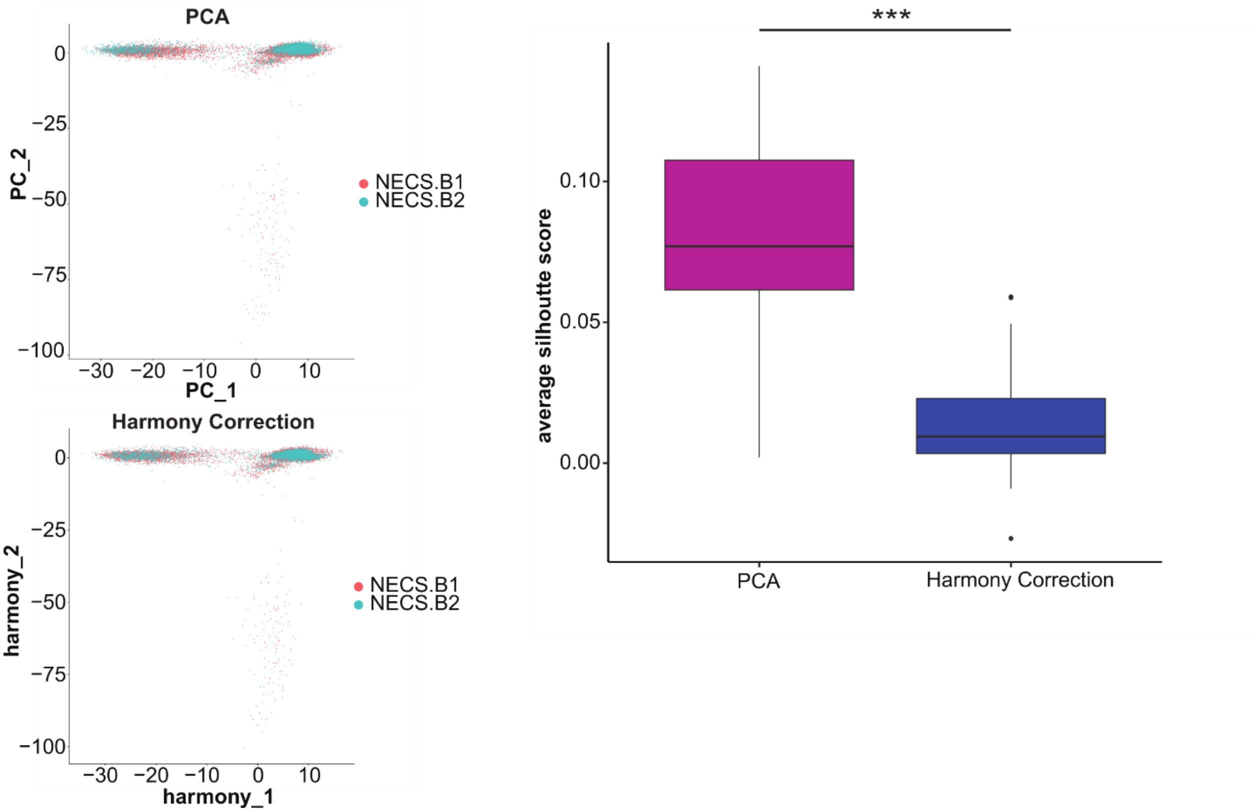


**Supplementary Figure 1. PCA correction for novel New England Centenarian single cell dataset. a,** Batch correction of novel dataset using Harmony based PCA correction: PCA plot before correction (top) and Harmony-corrected PCA (bottom). **b,** Average silhouette score for each immune subpopulation before and after Harmony correction to assess mixing of batches (T-test, p-value = 4.569e-06). A two-tailed T-test was performed with p-value significance of 0.05: nsp > 0.05, *p<0.05, **p<0.01, ***p<0.001.

A

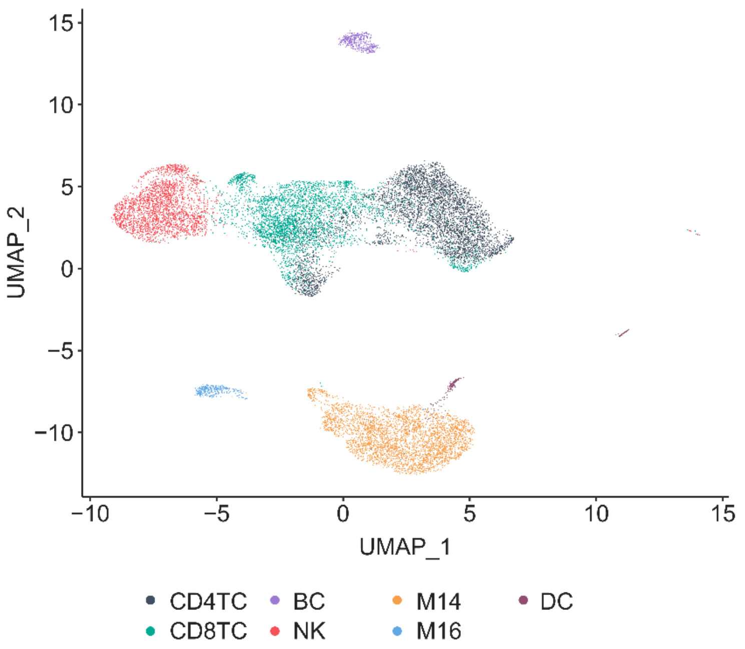


B

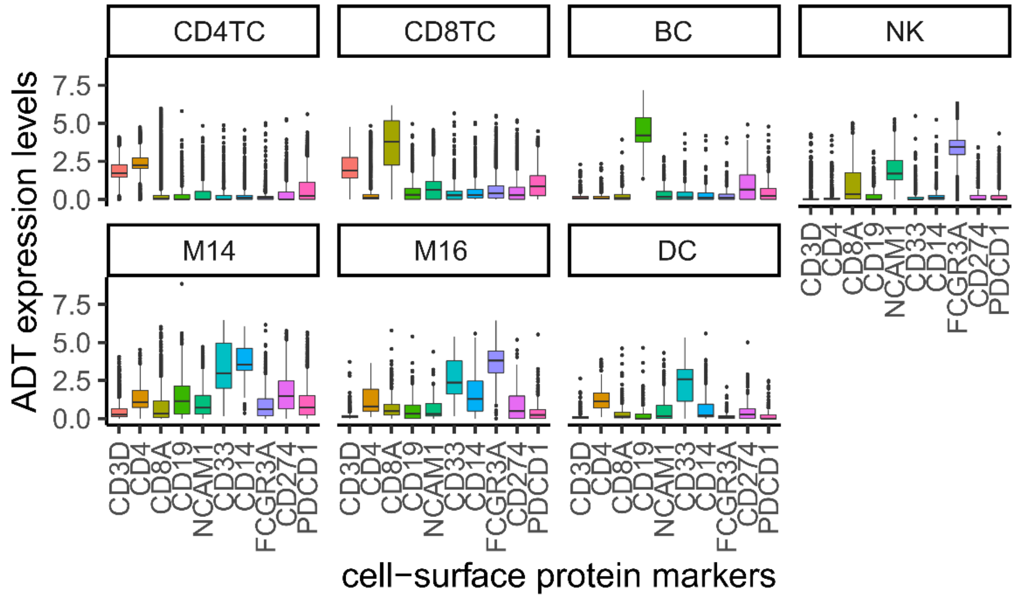


**Supplementary Figure 2. Protein expression levels across main immune populations of novel NECS dataset.** Boxplots of normalized protein expression levels (y-axis) of 10 protein markers (x-axis) across immune populations of novel NECS datasets: CD4 T cells (CD4TC), CD8 T cells (CD8TC), Natural Killer cells (NK), B cells (BC), Monocytes (Mono: M14 and M16), and Dendritic cells (DC).

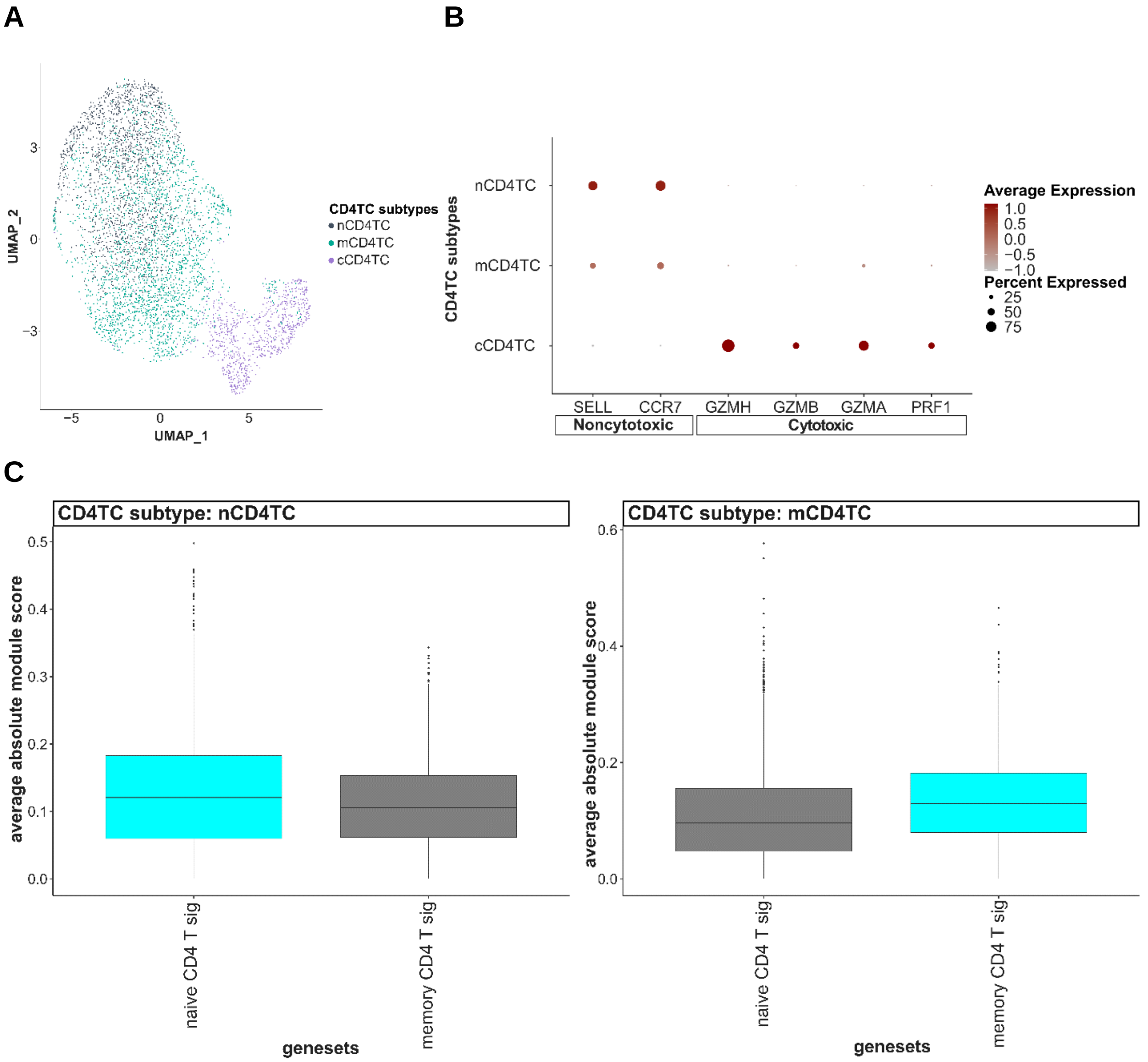


**Supplementary Figure 3. Identification of CD4TC subtypes in novel NECS dataset.** **a)** UMAP plot of CD4TC clustered and identified by subtypes: nCD4TC, mCD4TC, and cCD4TC. **b)** Dotplot of average expression of noncytotoxic-specific genes (SELL, CCR7) and cytotoxic specific genes (GZMH, GZMB, GZMA, PRF1) in CD4TC subtypes: nCD4TC, mCD4TC, and cCD4TC. **c)** Boxplots of average absolute module scores (y-axis) of naive and memory CD4 T cell signatures (x-axis) for each CD4TC subtypes from NECS datasets: nCD4TC and mCD4TC.

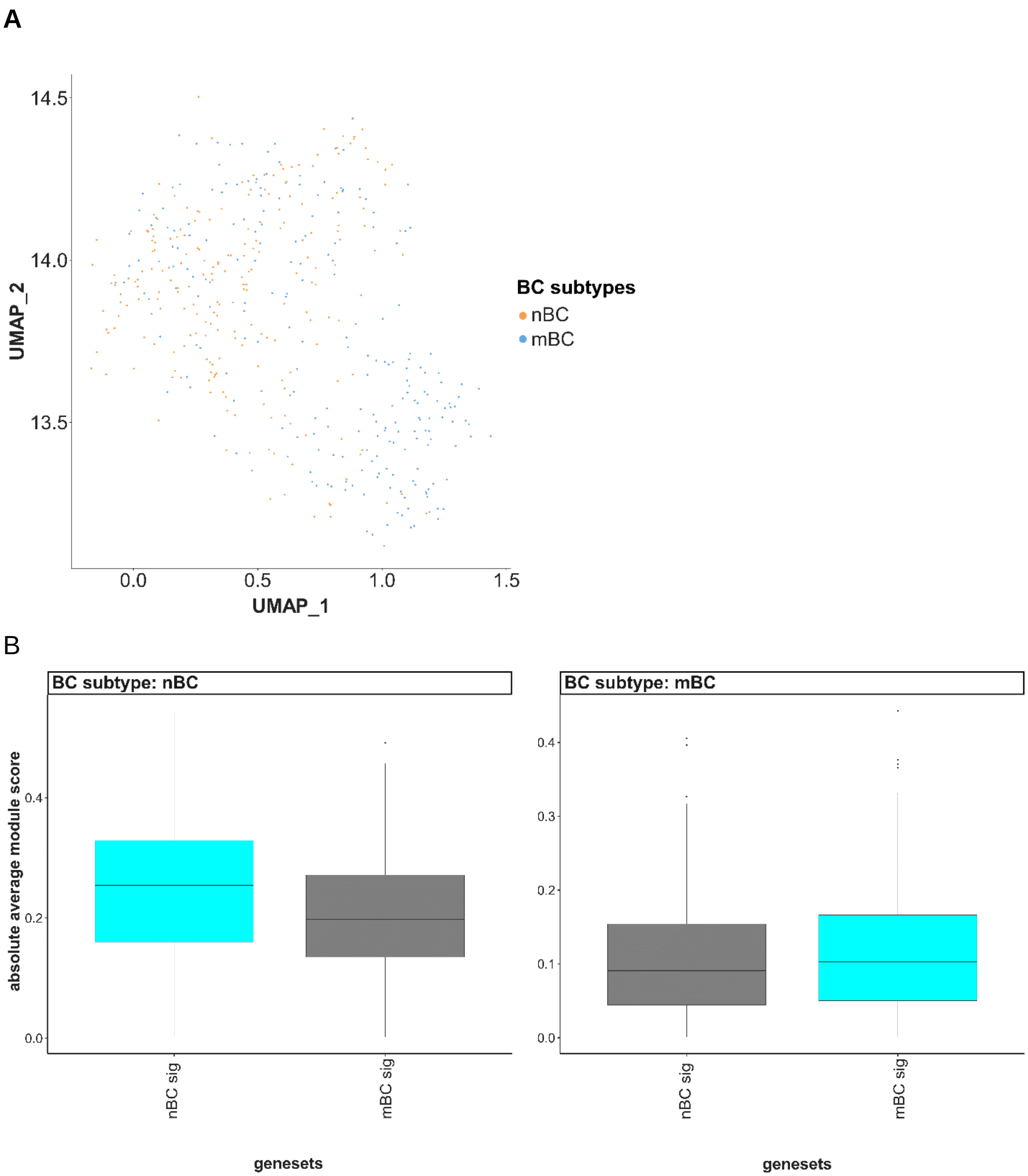


**Supplementary Figure 4. Identification of BC subtypes in novel NECS dataset. a)** UMAP plot of BC clustered and identified by subtypes: nBC and mBC. **b)** Boxplots of average absolute module scores (y-axis) of nBC and mBC signatures (x-axis) for each BC subtype from NECS datasets: nBC and mBC.

A

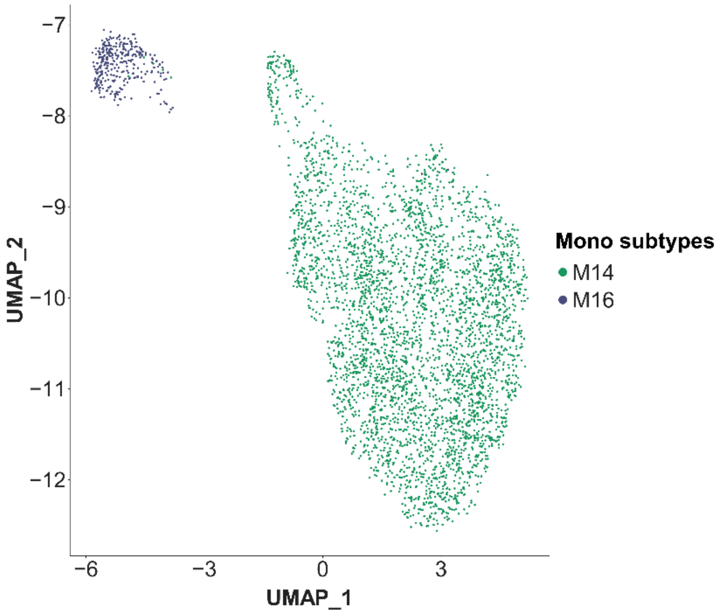


B

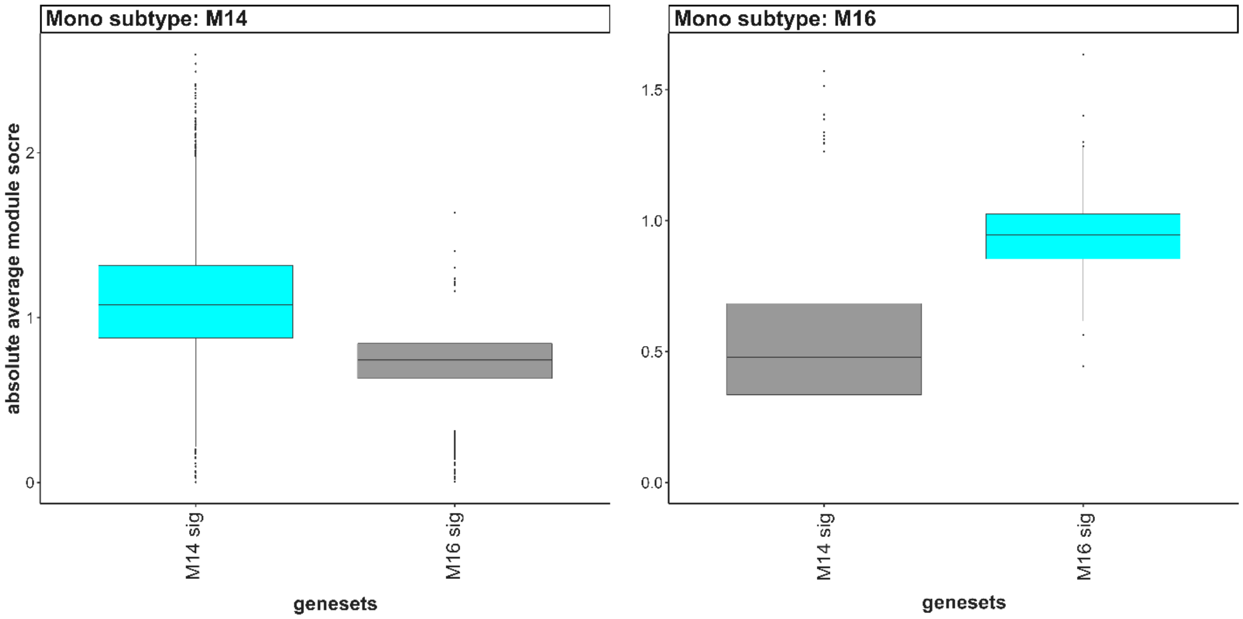


**Supplementary Figure 5. Identification of Monocyte subtypes in novel NECS dataset. a)** UMAP plot of Mono clustered and identified by subtypes: M14 and M16**. b)** Boxplots of average absolute module scores (y-axis) of M14 and M16 signatures (x-axis) for each Mono subtype from NECS datasets: M14 and M16.

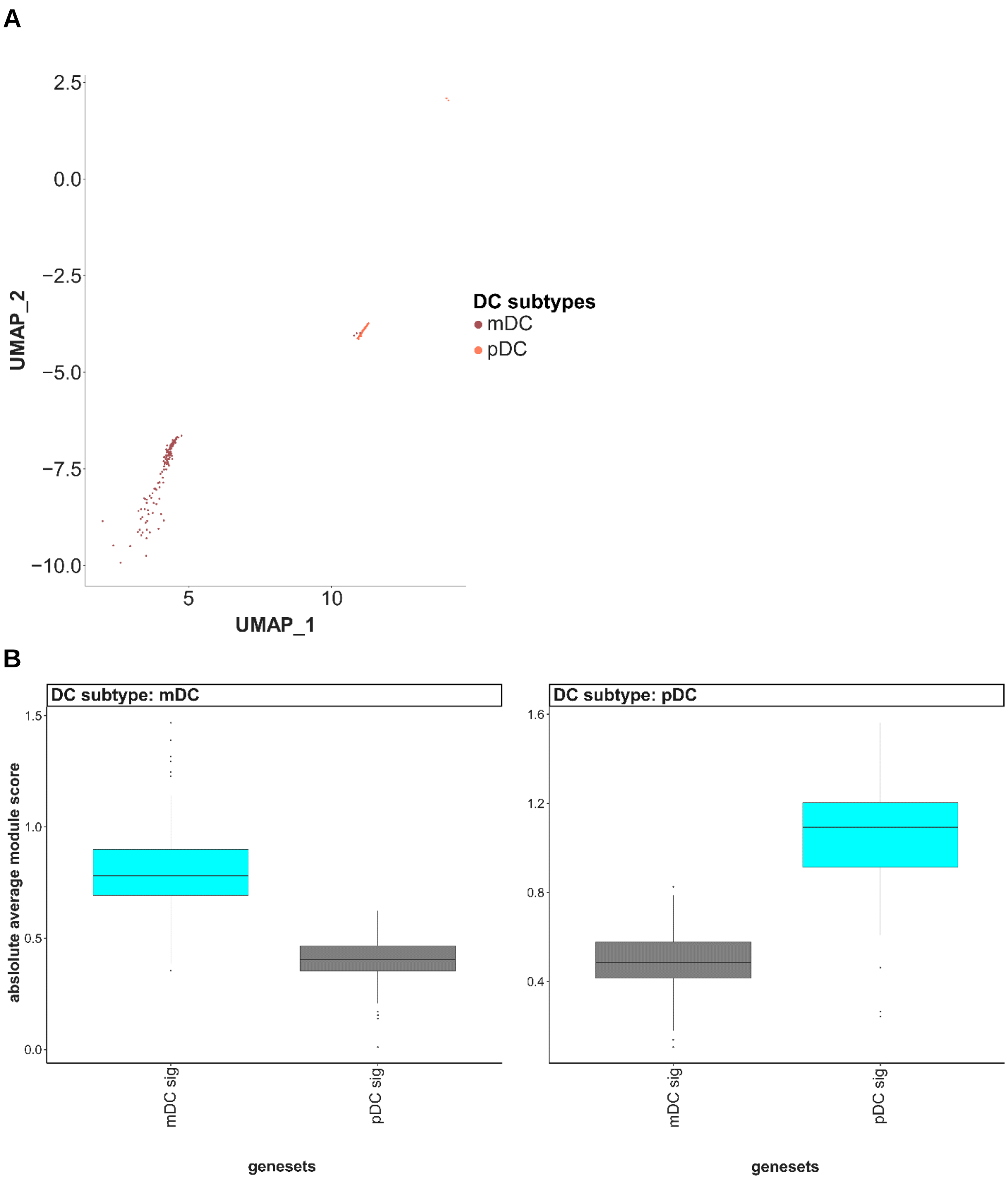


**Supplementary Figure 6. Identification of DC subtypes in novel NECS dataset. a)** UMAP plot of DC clustered and identified by subtypes: mDC and pDC **. b)** Boxplots of average absolute module scores (y-axis) of mDC and pDC signatures (x-axis) for each DC subtype from NECS datasets:  mDC and pDC.

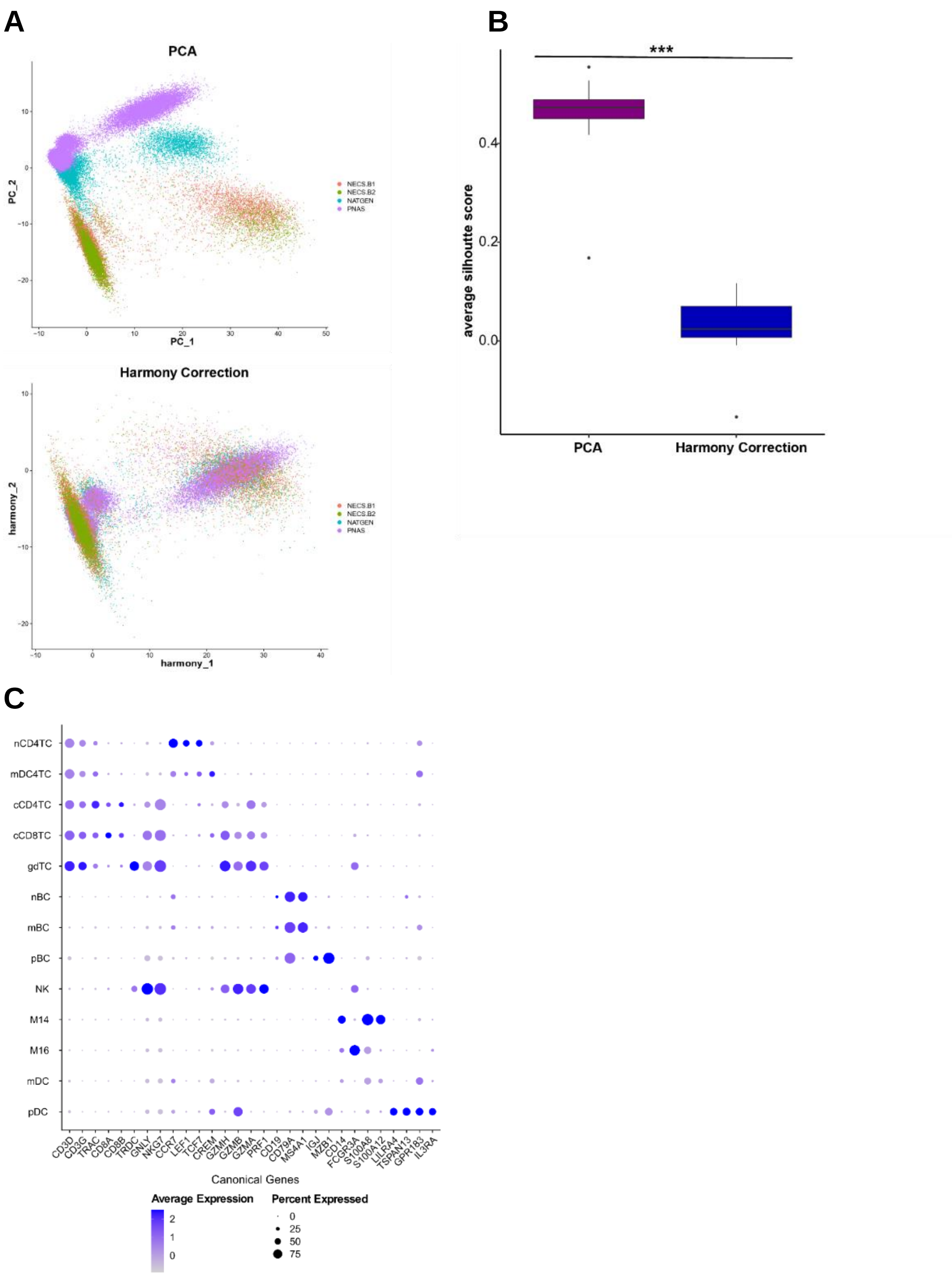


**Supplementary Figure 7. Harmonization of NECS and publicly available single cell datasets of aging and longevity**. **a,** Batch correction of NECS with PNAS and NATGEN datasets using Harmony based PCA correction: PCA plot before correction (top) and Harmony-corrected PCA. **b,** Average silhouette score for each immune subpopulation before and after Harmony correction to assess mixing of datasets (T-test, p-value = 1.518e-08). A two-tailed T-test was performed with p-value significance of 0.05: nsp > 0.05, *p<0.05, **p<0.01, ***p<0.001. **c,** Dot plot of average expression of canonical gene markers of immune cell populations in the integrated datasets.

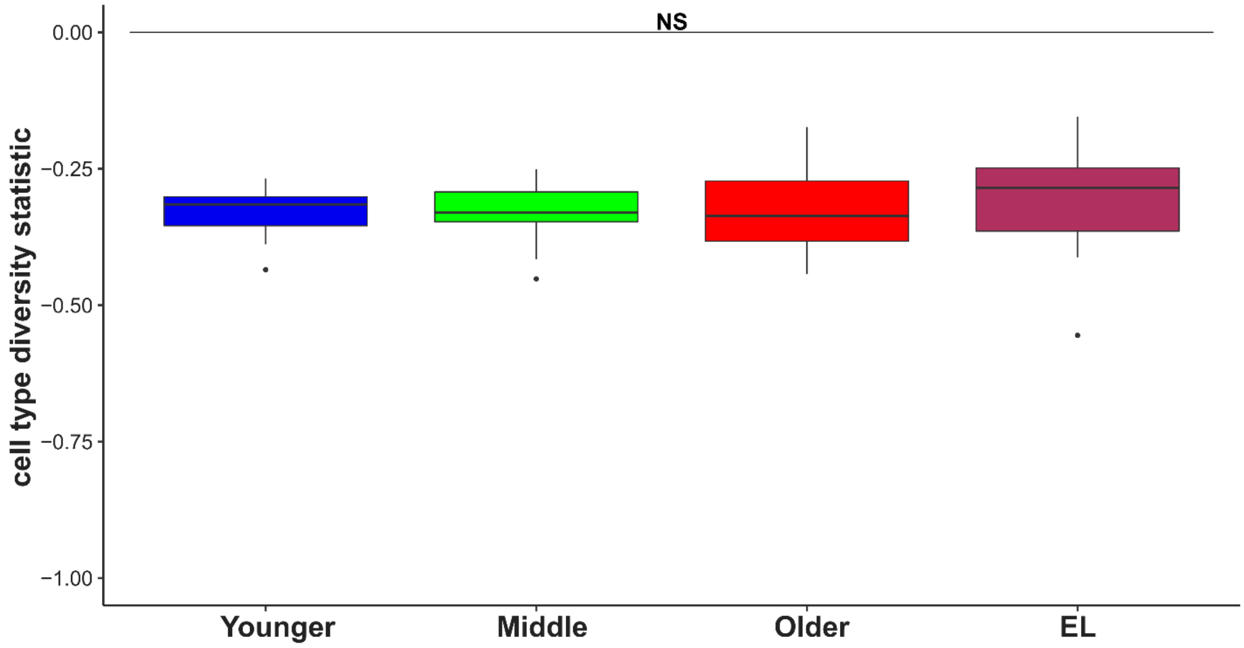


**Supplementary Figure 8. Cell type diversity statistic based on the 13 immune subtypes.** Boxplot of cell type diversity statistic calculated on the 13 immune cell subtypes per sample and grouped across age groups, with increased cell type diversity in EL compared to younger age groups although not found to be statistically significant (F-test, p-value = 0.7231).

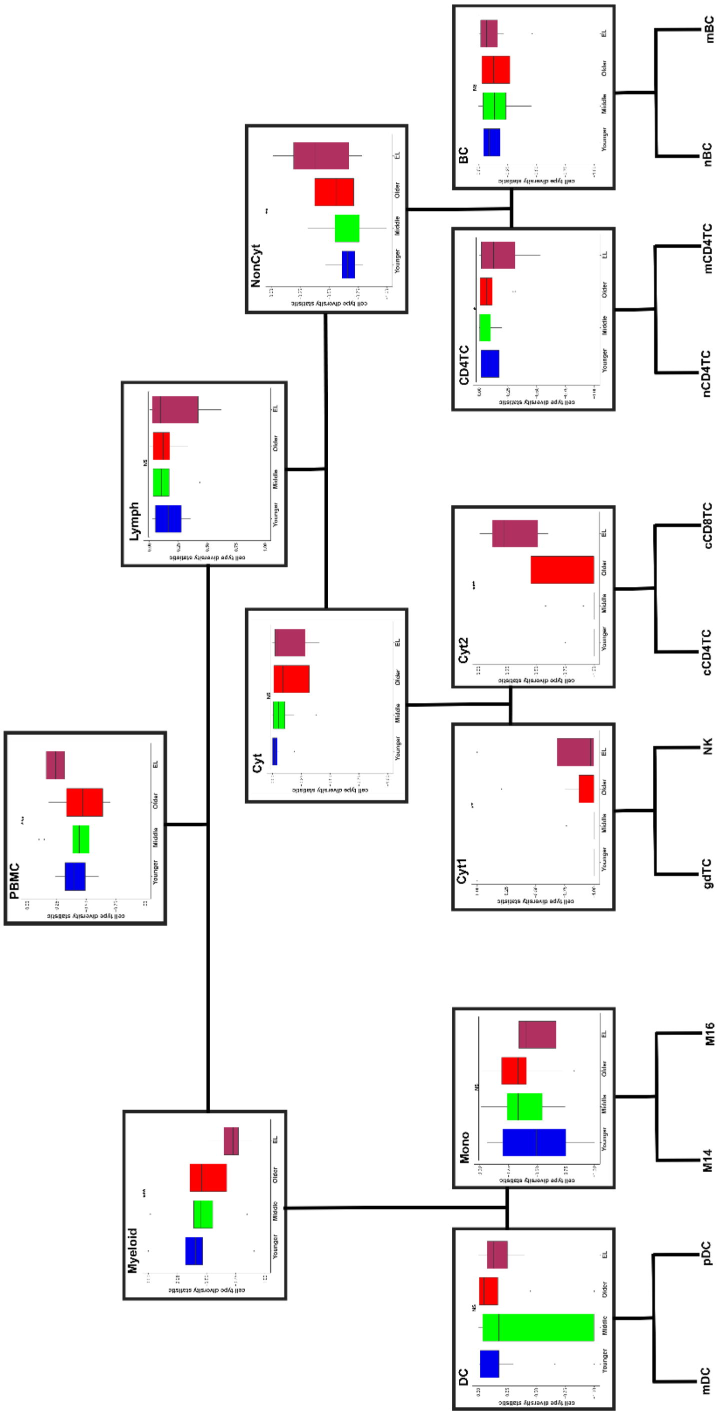


**Supplementary Figure 9. Diversity along the hierarchy of peripheral immune compartments.** Boxplots of the cell type diversity statistic for each sample based on the cell type proportions within each peripheral immune compartment compared across age groups.

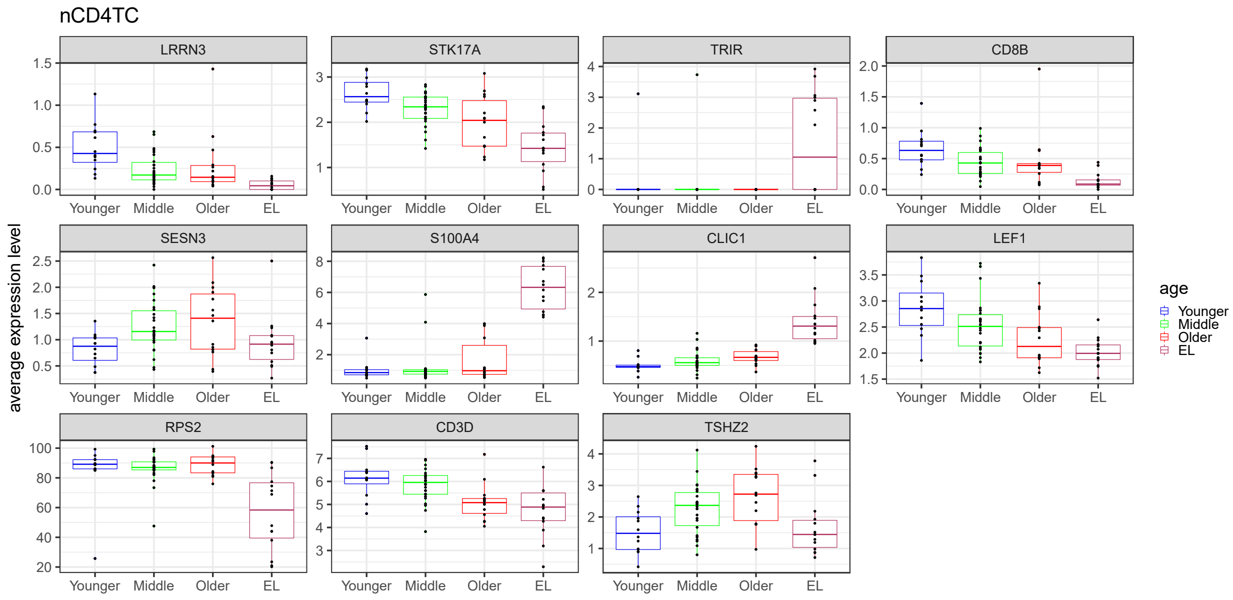


**Supplementary Figure 10. Transcriptional signatures across the human lifespan in nCD4TC.** Boxplots of average expression levels per sample of significant differential genes in nCD4TC across younger age, middle age, older age, and EL displaying age-related patterns.

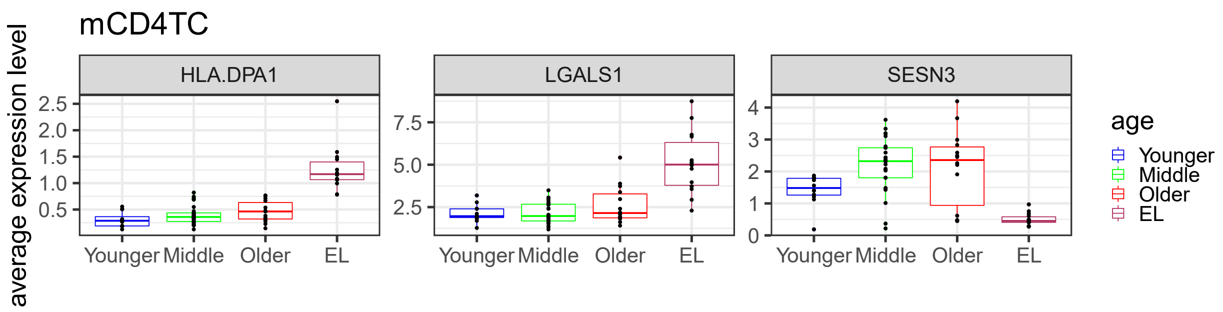


**Supplementary Figure 11. Transcriptional signatures across the human lifespan in mCD4TC.** Boxplots of average expression levels per sample of significant differential genes in mCD4TC across younger age, middle age, older age, and EL displaying age-related patterns.

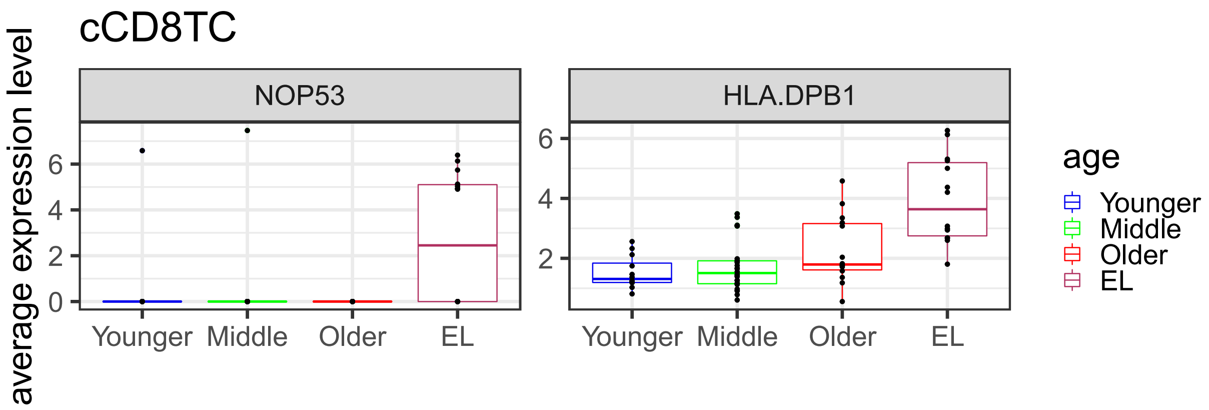


**Supplementary Figure 12. Transcriptional signatures across the human lifespan in cCD8TC.** Boxplots of average expression levels per sample of significant differential genes in cCD8TC across younger age, middle age, older age, and EL displaying age-related patterns.

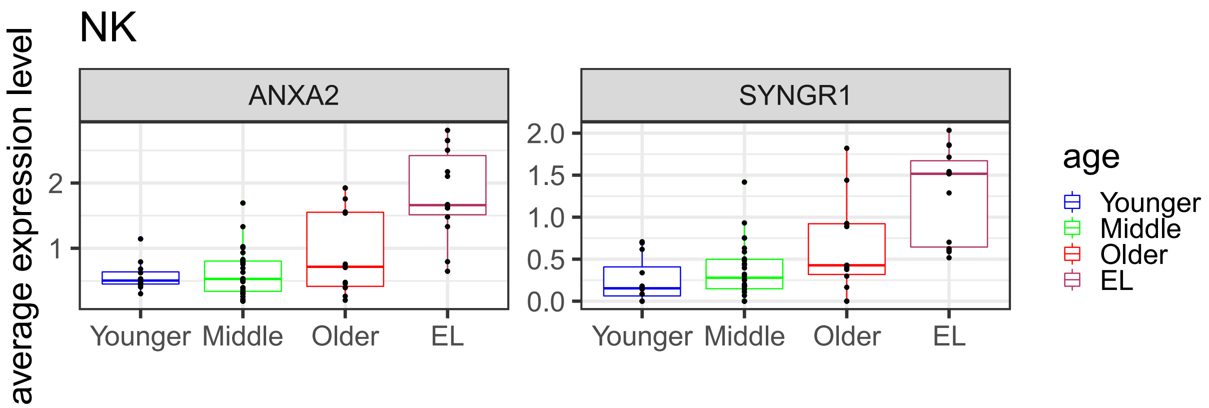


**Supplementary Figure 13. Transcriptional signatures across the human lifespan in NK.** Boxplots of average expression levels per sample of significant differential genes in NK across younger age, middle age, older age, and EL displaying age-related patterns.

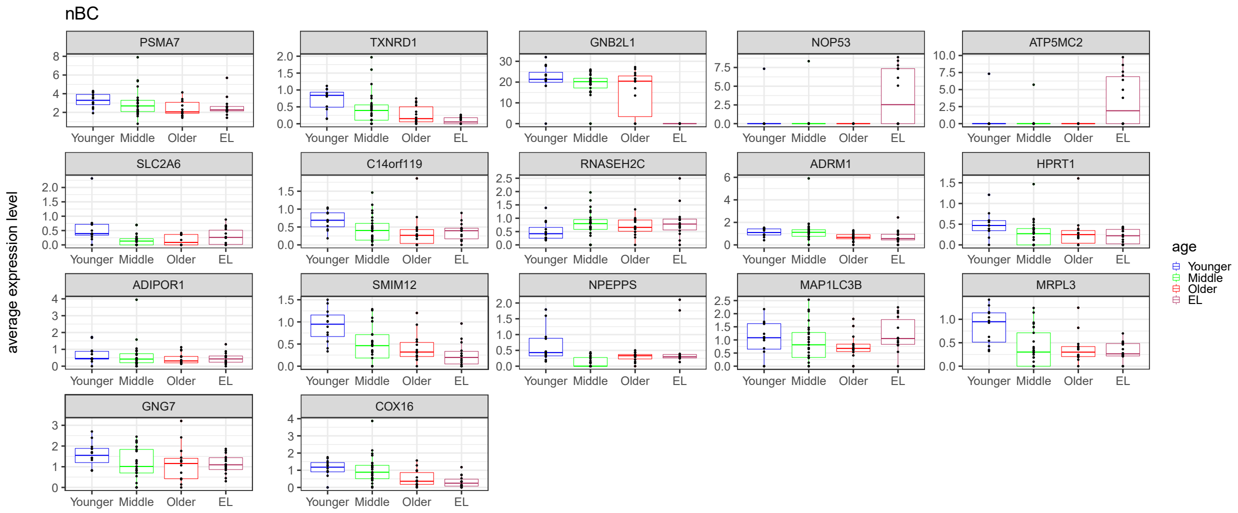


**Supplementary Figure 14. Transcriptional signatures across the human lifespan in nBC.** Boxplots of average expression levels per sample of significant differential genes in nBC across younger age, middle age, older age, and EL displaying age-related patterns.

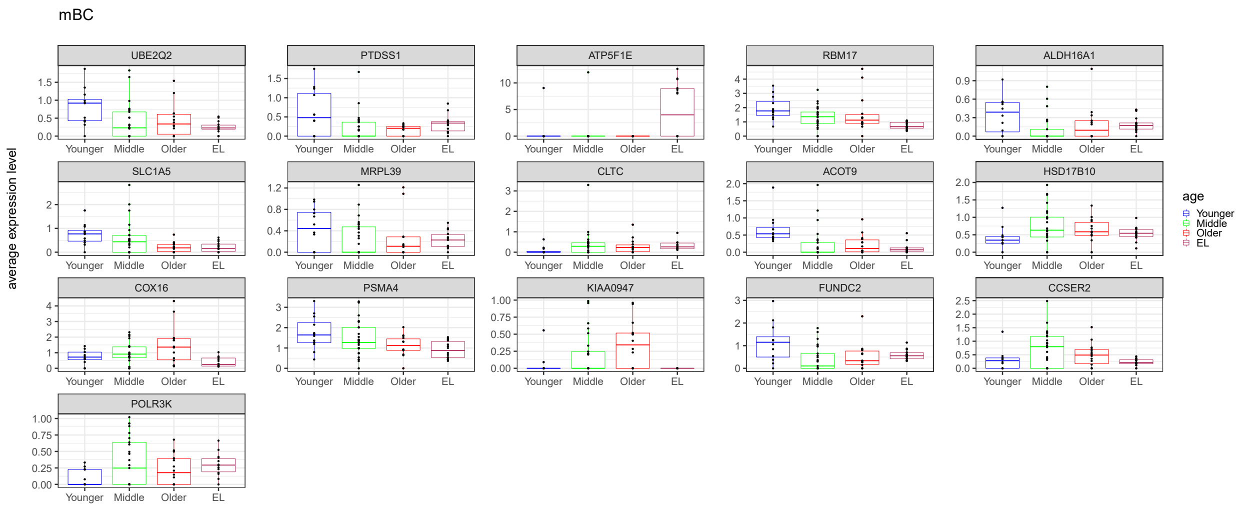


**Supplementary Figure 15. Transcriptional signatures across the human lifespan in mBC.** Boxplots of average expression levels per sample of significant differential genes in mBC across younger age, middle age, older age, and EL displaying age-related patterns.

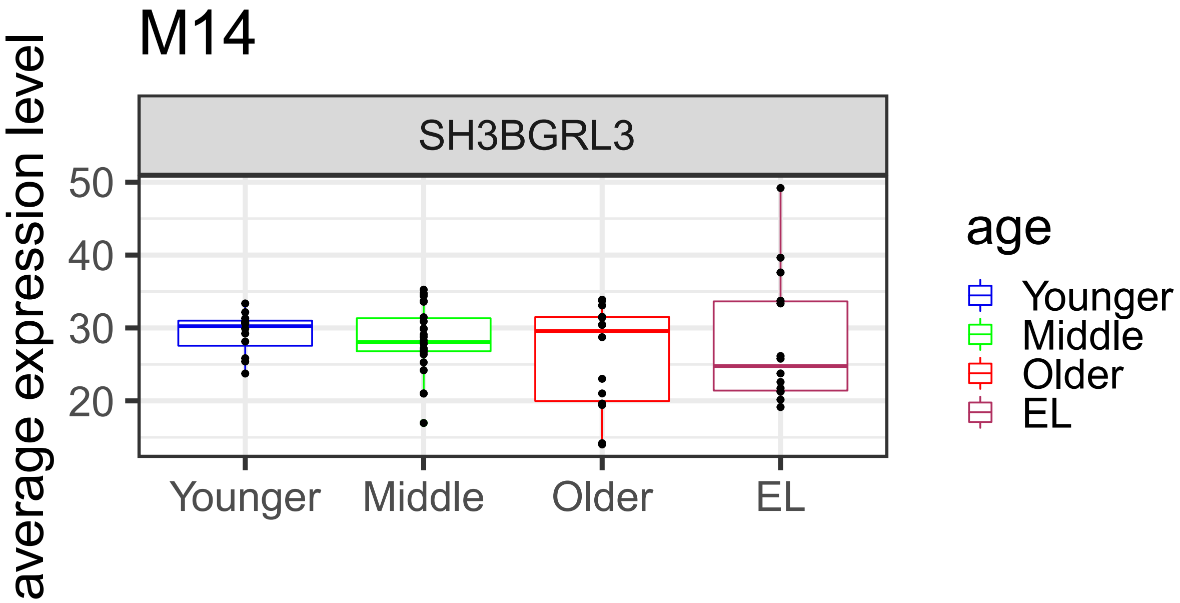


**Supplementary Figure 16. Transcriptional signatures across the human lifespan in M14.** Boxplots of average expression levels per sample of significant differential genes in M14 across younger age, middle age, older age, and EL demonstrating patterns of extreme old age.

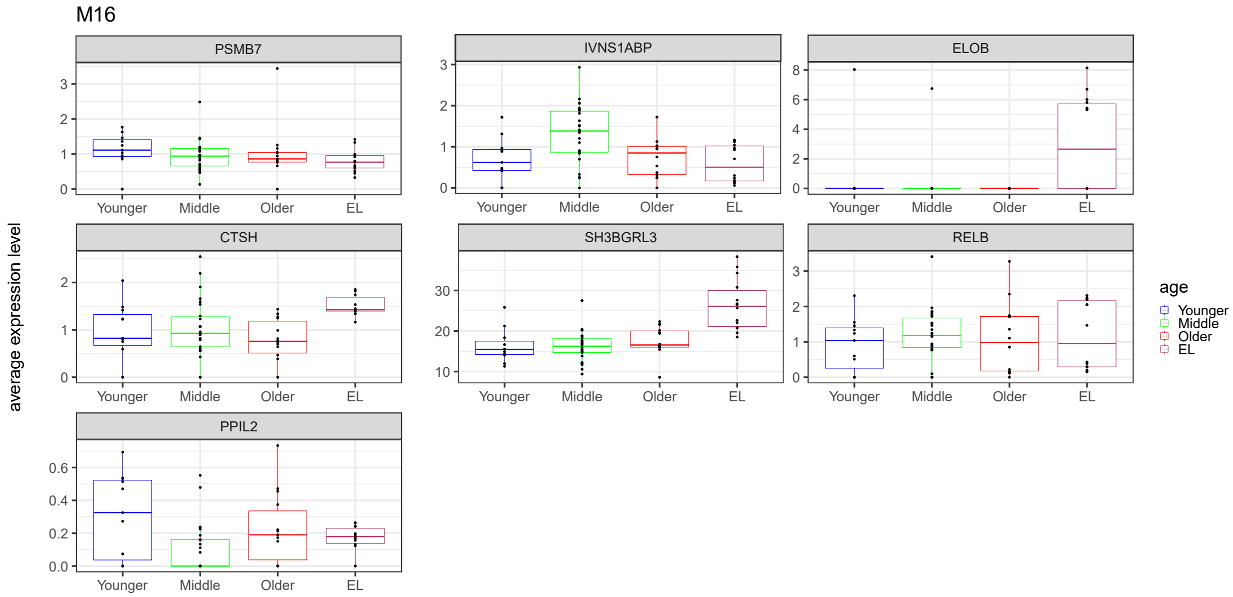


**Supplementary Figure 17. Transcriptional signatures across the human lifespan in M16.** Boxplots of average expression levels per sample of significant differential genes in M16 across younger age, middle age, older age, and EL demonstrating patterns of extreme old age.

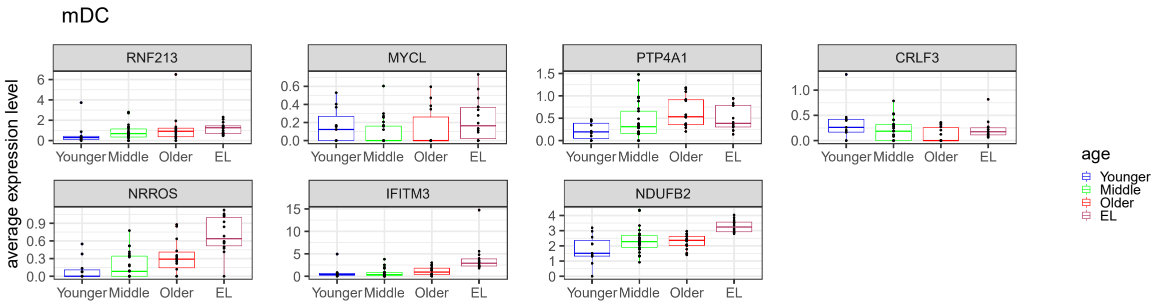


**Supplementary Figure 18. Transcriptional signatures across the human lifespan in mDC.** Boxplots of average expression levels per sample of significant differential genes in mDC across younger age, middle age, older age, and EL demonstrating patterns of extreme old age.

A

| | Middle v. Younger | Older v. Younger | EL v. Younger |
|---|---|---|---|
| **Upregulated** | **0** | **1** | **136** |
| **Downregulated** | **0** | **2** | **251** |

B

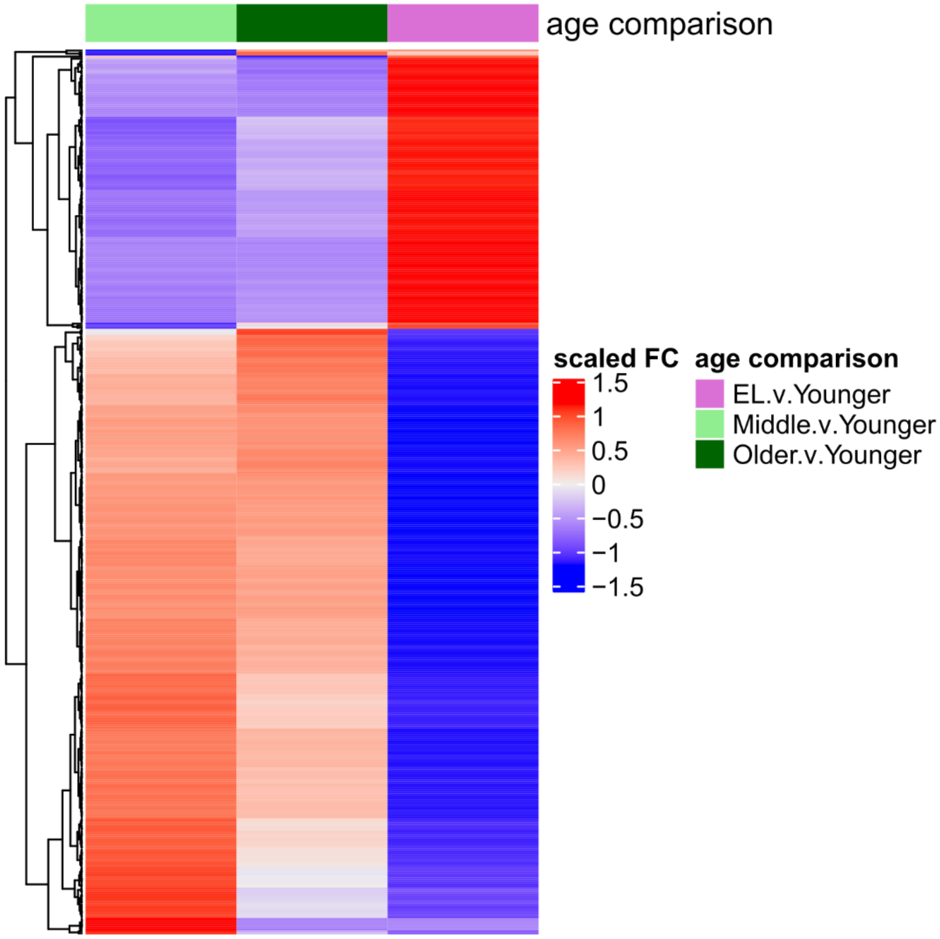


**Supplementary Figure 19. Bulk level gene expression changes across the human life span. a,** Table of number of significant genes upregulated and downregulated for each age comparison: Middle v. Younger age, Older v. Younger age, and EL v. Younger age. **b,** Heatmap of bulk level significant genes across sample level expression data.
